# A Candidate Trajectory-Atypicality Marker of Concurrent Cognitive Change in Alzheimer’s Disease from a Spatio-Temporal Graph Attention Network

**DOI:** 10.64898/2026.11.25.769046

**Authors:** Arman Dinarvand, Amirali Moradniaei, Elaheh Amini, Keivan Borna, the Alzheimer’s Disease Neuroimaging Initiative

## Abstract

Structural MRI markers of Alzheimer’s disease (AD) quantify tissue loss, yet patients with similar atrophy follow different clinical courses. We trained a spatio-temporal graph attention network on longitudinal morphometry of five executive-control and salience-network regions and derived an off-manifold residual that measures how atypical a participant’s trajectory is. In 305 participants from the Alzheimer’s Disease Neuroimaging Initiative (ADNI-1), each embedded by a fold-specific network that was not trained on them, the residual was higher in AD than in cognitively normal participants and participants with mild cognitive impairment (Kruskal–Wallis *H* = 20.8, *ε*^2^ = 0.068, *p <* 0.001) and was nominally associated with atrophy (Spearman *ρ* = 0.12, *p* = 0.035), an association that did not survive correction across four covariates (*q* = 0.14). Six cognitive endpoints were tested without a pre-registered hierarchy and corrected together. In a prospective model given only baseline and month-12 imaging, the association of the residual with 24-month change in the Mini-Mental State Examination, adjusted for atrophy and baseline cognition, was nominal and did not survive this correction (partial *ρ* = *−*0.14, *p* = 0.017, *q* = 0.10). With whole acquisition sites withheld from training it did (*ρ* = *−*0.20, *q* = 0.003), but the two estimates did not differ significantly (difference *−*0.06, 95% CI *−*0.14 to 0.02). No other endpoint was associated with the residual after correction. The evidence was limited in four further respects. The 24-month change overlaps the month-12 imaging input, and neither prospective model was associated with change from month 12 to month 24 (partial *ρ* = *−*0.05 and *−*0.04, *p ≥* 0.42). The association was not significant within AD (*ρ* = *−*0.12, *p* = 0.27). Adding the residual to atrophy increased the area under the receiver operating characteristic curve for AD by 0.030 to 0.041, but every 95% confidence interval for the increase included zero. The association was reduced to near zero when supervision of the regional atrophy rate was removed during training (*ρ* = *−*0.03, *p* = 0.56), and the same ablation in the site-disjoint folds also removed it (*ρ* = *−*0.09, *p* = 0.13). The atypicality was distributed across salience and executive-control regions. The residual is a candidate marker of cognitive change over the imaging window; whether it predicts decline after that window requires independent validation.

## 1 Introduction

Alzheimer’s disease (AD) unfolds over years, and its rate of progression varies considerably between individuals [1–3]. Most imaging biomarkers report the damage a brain has already sustained, such as hippocampal volume, cortical thickness, or a regional atrophy score. These measures are informative but static: two patients with the same volumetric score can have very different futures, depending on whether their decline is stabilizing or accelerating. Clinical management and trial design would benefit from a measure of how a brain network is moving through its disease course, and of whether that motion is typical for the disease.

Characterizing progression requires both the spatial relationships among brain regions and the temporal changes in regional morphology. Interregional relationships carry information beyond individual regional measurements [4–6], and longitudinal data capture the rate and curvature of change, which cannot be estimated from a single scan [7, 8]. Cross-sectional classifiers do not model within-person trajectories, and many deep-learning pipelines for AD are trained and evaluated at a single visit [9]; region-wise longitudinal regression, in turn, does not explicitly represent interregional relationships.

Graph neural networks address the spatial component. Graph convolutional networks introduced learned neighbourhood aggregation on irregular topologies [10], and attention-based variants, GAT [11] and GATv2 [12], learn which connections matter rather than fixing them in advance. In neuroimaging, BrainNetCNN [13], BrainGNN [14], and the Brain Network Transformer [15] build on this idea and on the broader success of attention [16], but almost always at one timepoint. Disease-progression models address the temporal component through continuous-time event models [17], subtype-and-stage inference [18], and the TADPOLE challenge [19], but treat regions independently or rely on hand-engineered staging variables.

The clinically relevant quantity is how the state changes, and whether the rate of change is itself changing. A model that consumes three visits can read out velocity and acceleration directly, so that its representation reflects trajectory shape rather than position; this motivates the temporal difference encoder used here. We model regions of the executive-control and salience networks rather than the medial temporal lobe. The salience network is the principal target of behavioural-variant frontotemporal dementia [4], and in AD its connectivity is first increased and then lost as dementia progresses, alongside loss within and between other large-scale networks [20, 21]. Together with the executive-control network, it underlies the executive and attentional deficits that accompany decline [22]. These networks are less established as structural-MRI markers of AD than the medial temporal lobe; a trajectory signal in these regions therefore tests a different system, and it is a harder test for a graph model, which must use network structure rather than large focal volume loss. *APOE ε*4 is the strongest common genetic risk factor for late-onset AD [23–25] and has non-uniform regional effects, which motivates conditioning the network on genotype.

Because the cohort is modest in size, we augmented the training data with synthetic participants from a tabular diffusion model and tested their similarity to the real participants before analysing the embeddings. Each participant’s embedding was generated by a fold-specific model that was not trained on that participant. Three configurations support the main analyses: a three-visit model, a two-visit (prospective) model that does not receive the month-24 scan, and a site-disjoint retrain of the prospective model; Table 2 lists these configurations together with their controls. The study makes five contributions. First, we introduce an off-manifold residual, a geometric measure of trajectory atypicality that is elevated in AD and only nominally associated with atrophy. Second, we show that the residual of the prospective model is associated with change on the Mini-Mental State Examination from baseline to month 24, an interval that includes its month-12 imaging input, when whole acquisition sites are withheld from training, whereas with a stratified split the association is nominal and does not survive correction across six cognitive endpoints; neither split showed an association with change from month 12 to month 24, after the last imaging input, so the residual tracks cognitive change over the imaging window rather than predicting later decline. Third, we show by ablation that this association depends on supervision of the regional atrophy rate rather than on an auxiliary tipping-point objective. Fourth, we localize the atypicality to salience and executive-control regions and to attention reconfiguration on the insula–parietal connection. Fifth, we show that a diffusion pseudo-time readout of the same embedding is a graph artifact, and we describe controls that distinguish such artifacts from true associations.

## 2 Methods

### 2.1 Participants

Data used in the preparation of this article were obtained from the Alzheimer’s Disease Neuroimaging Initiative (ADNI) database (adni.loni.usc.edu). The ADNI was launched in 2003 as a public-private partnership, led by Principal Investigator Michael W. Weiner, MD. The primary goal of ADNI has been to test whether serial magnetic resonance imaging (MRI), positron emission tomography (PET), other biological markers, and clinical and neuropsychological assessment can be combined to measure the progression of mild cognitive impairment (MCI) and early Alzheimer’s disease (AD).

Participants were drawn from the ADNI-1 phase. No formal sample-size calculation was performed; all participants who met the following criteria were analysed. We required structural MRI at baseline, month 12, and month 24, baseline *APOE* genotyping, and a clinical diagnosis at each visit in one of the ADNI categories (cognitively normal, CN; MCI; AD) [1]. We excluded participants with mid-study diagnostic reversal, cortical segmentations that failed visual quality control at any visit, or more than 25% missing values across the derived regional features. The final cohort comprised 305 participants (Table 1). Two AD participants whose month-24 MMSE was recorded with an ADNI missing-value code were excluded from analyses of MMSE change, which include 303 participants (89 AD).

**Table 1.** Cohort characteristics. Mean (standard deviation) or *n* (%) by baseline diagnosis. *p*: Kruskal–Wallis for continuous variables and *χ*^2^ for categorical variables (for *APOE*, across zero, one, and two *ε*4 alleles). MMSE change is the month-24 minus the baseline score; negative values indicate decline; it is available for 303 participants (89 AD). CN, cognitively normal; MCI, mild cognitive impairment; AD, Alzheimer’s disease. Education was not extracted for this analysis.

|  | CN | MCI | AD | All | *p* |
|---|---|---|---|---|---|
| Participants, *n* | 85 | 129 | 91 | 305 |  |
| Age, years | 76.2 (4.8) | 74.0 (7.0) | 76.1 (7.7) | 75.2 (6.7) | 0.051 |
| Female, *n* (%) | 49 (58%) | 50 (39%) | 43 (47%) | 142 (47%) | 0.025 |
| *APOE* *ε*4 alleles, *n* (%) |  |  |  |  | < 10<sup>−7</sup> |
| 0 | 63 (74%) | 57 (44%) | 27 (30%) | 147 (48%) |  |
| 1 | 20 (24%) | 56 (43%) | 45 (49%) | 121 (40%) |  |
| 2 | 2 (2%) | 16 (12%) | 19 (21%) | 37 (12%) |  |
| Baseline MMSE | 29.2 (0.9) | 27.3 (1.7) | 23.4 (2.1) | 26.6 (2.8) | < 10<sup>−40</sup> |
| 24-month MMSE change | −0.1 (1.1) | −1.7 (3.4) | −4.6 (5.7) | −2.1 (4.2) | < 10<sup>−10</sup> |

**Table 2.**
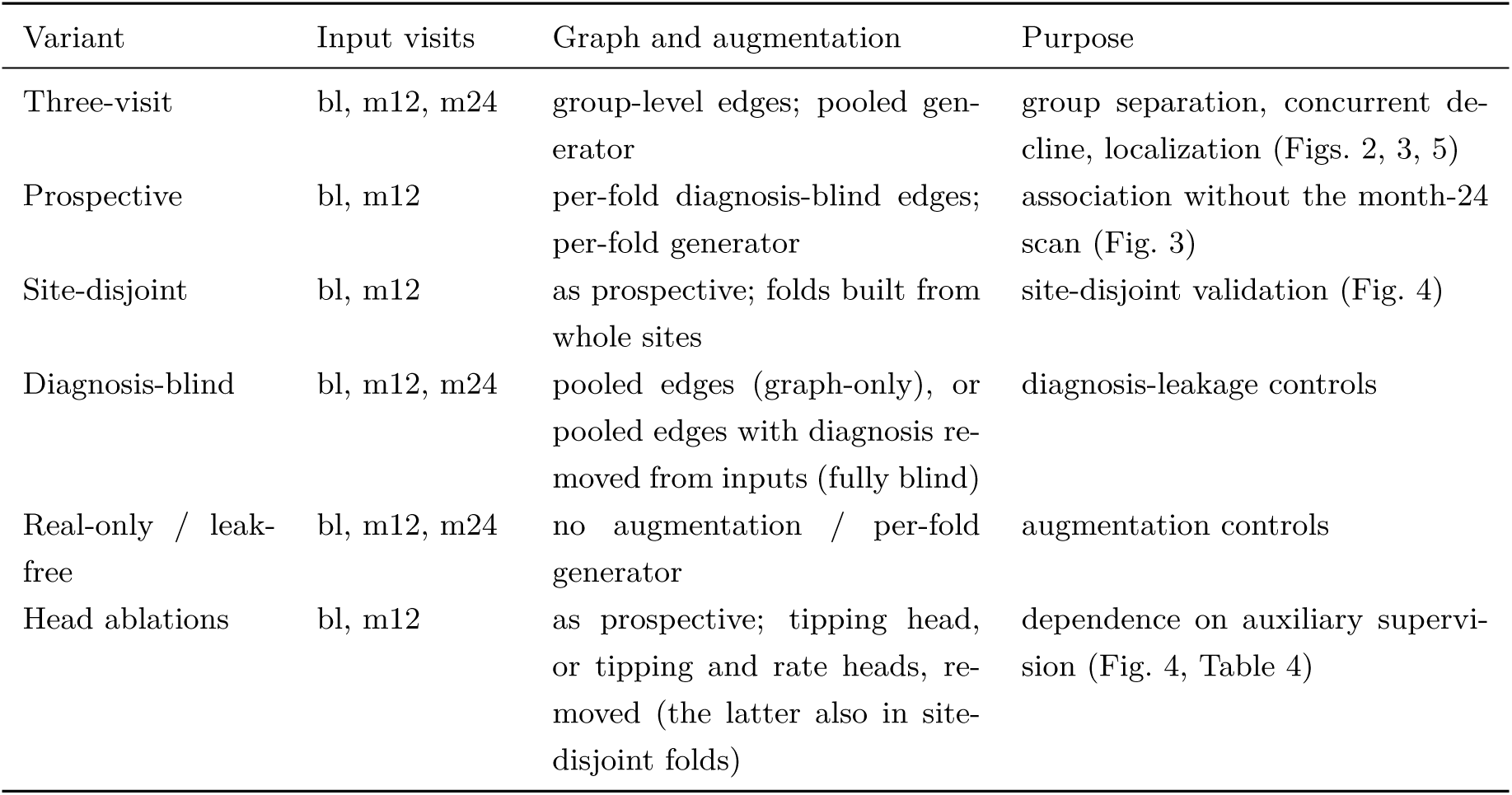
Model configurations and controls. The configurations share the architecture in Fig. 1 but differ in input visits, diagnostic metadata, graph construction, augmentation, and auxiliary heads as indicated. The diagnosis-blind, augmentation-control, and head-ablation rows each include two configurations. Each participant’s embedding is generated by a fold-specific model that was not trained on that participant, and residuals are computed within fold. The first three rows describe the principal analyses. bl, baseline; m12, month 12; m24, month 24.

| Variant | Input visits | Graph and augmentation | Purpose |
|---|---|---|---|
| Three-visit | bl, m12, m24 | group-level edges; pooled gen-erator | group separation, concurrent de-cline, localization (Figs. 2, 3, 5) |
| Prospective | bl, m12 | per-fold diagnosis-blind edges; per-fold generator | association without the month-24 scan (Fig. 3) |
| Site-disjoint | bl, m12 | as prospective; folds built from whole sites | site-disjoint validation (Fig. 4) |
| Diagnosis-blind | bl, m12, m24 | pooled edges (graph-only), or pooled edges with diagnosis re-moved from inputs (fully blind) | diagnosis-leakage controls |
| Real-only / leak-free | bl, m12, m24 | no augmentation / per-fold generator | augmentation controls |
| Head ablations | bl, m12 | as prospective; tipping head, or tipping and rate heads, re-moved (the latter also in site-disjoint folds) | dependence on auxiliary supervi-sion (Fig. 4, Table 4) |

### 2.2 Feature extraction

T1-weighted scans were skull-stripped with deepmriprep [26], resampled to 1 mm isotropic resolution with FSL FLIRT [27], and processed with DL+DiReCT [28], which segments the cortex into the Desikan–Killiany parcellation [29] with a deep network and estimates cortical thickness by diffeomorphic registration. Regional volume and mean cortical thickness were taken from the per-region DL+DiReCT outputs. Five bilateral nodes were formed from Desikan–Killiany regions for their roles in executive control and salience processing [22, 30]: dorsolateral prefrontal cortex (dlPFC; rostral and caudal middle frontal), ventrolateral prefrontal cortex (vlPFC; pars opercularis and pars triangularis), dorsal anterior insula (dAI; insula), dorsal anterior cingulate cortex (dACC; caudal and rostral anterior cingulate), and superior parietal lobule (SPL; superior parietal). Node volume was the sum of the constituent regions’ volumes, and node thickness their volume-weighted mean thickness. The parcellation does not subdivide the insula or separate dorsal from pregenual cingulate cortex, so the dAI node is the whole insula and the dACC node the whole anterior cingulate; we keep these labels to match the network literature. Each region carried seven features per visit: volume, thickness, the first differences of volume and thickness from the preceding visit, expanding-window longitudinal rates of volume and thickness, and a volumetric asymmetry index (*L − R*)*/*(*L* + *R*). Rates were computed causally as the least-squares slope over visits up to and including the current one, so no feature contains information from a later visit. Features were z-scored per channel on the pooled cohort.

### 2.3 Graph construction

We constructed a complete directed graph with five nodes and 20 non-self edges; the attention layer added self-loops. Within each diagnostic group, edge weights were the Fisher z-transformed [31] Pearson correlations across participants between the bilateral baseline volumes of each pair of regions (structural covariance). The same weights were used at all three visits, so temporal information was represented by the encoder rather than by changes in the graph. For the diagnosis-blind graph, the correlations were computed across all participants without stratification by diagnosis.

### 2.4 Model architecture and training

A metadata projector maps a four-dimensional vector (*APOE* risk score, carrier flag, normalized risk score, and baseline diagnostic group) to 16 dimensions through two linear layers with layer normalization [32] and GELU activations; the projected context is concatenated to each node’s seven features. The spatial layer is a GATv2 convolution [12] with four heads, 32 hidden units per head, attention dropout 0.3, self-loops, and a one-dimensional edge attribute, implemented in PyTorch [33] with PyTorch Geometric [34]; head outputs are concatenated to 128 dimensions and layer-normalized. The temporal difference encoder takes the three spatial representations *h*^(1)^*, h*^(2)^*, h*^(3)^ and computes two first-order differences, *Δ^1/12^ = h^(2)^ − h^(1)^*, and one second-order difference, *Δ^2^ = Δ^1/23^ − Δ^1/12^*. Each difference is projected to 64 dimensions by a separate two-layer multilayer perceptron (MLP), and the projected differences are combined with the baseline representation into a 64-dimensional embedding *h*_node_ for each node (Fig. 1). Linear layers use Xavier initialization, and the model has approximately 2 *×* 10^5^ trainable parameters.

**Figure 1.**
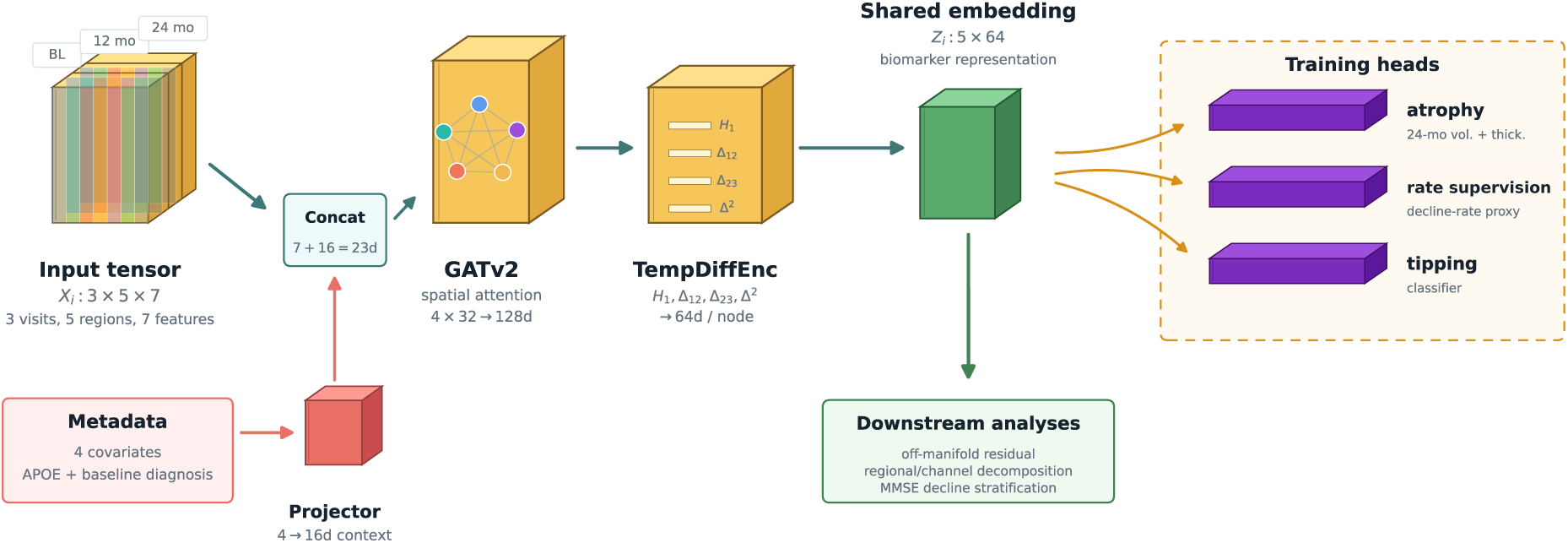
Model architecture. Each participant contributes three visits of five executive-control and salience regions with seven morphometric features per region (3 *×* 5 *×* 7) and four metadata covariates, which are projected to a 16-dimensional context vector and concatenated to every node. A GATv2 layer (four heads, 32 units per head) produces a 128-dimensional representation per node and visit, and a temporal difference encoder combines the baseline state, first differences, and second difference into a 64-dimensional node embedding (5 *×* 64 per participant). Three task heads (atrophy, rate supervision, tipping point) share this embedding during training. The residual-based analyses use the embedding; the supplementary uncertainty and attribution analyses also use the atrophy-head output.

Three heads share the embedding during training: an atrophy head predicting month-24 regional volume and thickness, a rate-supervision head predicting the negative z-scored month-12 regional volume-decline rate, and a tipping-point classifier. The heads shape the shared embedding during training, and their outputs are not used as biomarkers. The tipping target is a near-deterministic function of within-subject volume slope and month-12 volume (Supplementary Note 5). Its loss therefore acts as an auxiliary regularizer of the representation, not as supervision from an independent clinical endpoint, and high tipping-head accuracy does not indicate prediction of a clinical tipping point. Its output is excluded from downstream biomarker analyses.

Task losses (mean squared error for the atrophy and rate targets, binary cross-entropy for the tipping target) were combined with uncertainty weighting [35],

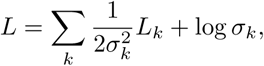

with a learned log-variance per task. Optimization used AdamW [36] (learning rate 10*^−^*^3^, weight decay 10*^−^*^4^) with cosine annealing to 10*^−^*^5^ over 100 epochs [37], gradient-norm clipping at 1.0, and early stopping with patience 15. Early stopping and checkpoint selection used the multitask morphometric loss on the held-out fold; this loss contains no cognitive score or diagnostic label, but it makes the held-out fold part of model selection. We therefore retrained the prospective and site-disjoint models with early stopping and checkpoint selection on an inner validation set of 15% of each fold’s real training participants (stratified by diagnosis), which were excluded from gradient updates, so that the held-out fold was not evaluated during training. Models were trained under five-fold cross-validation stratified by baseline diagnosis. Each configuration was trained once, with random seed 42 (42 plus the fold index for per-fold generators and prospective retrains); no repeated-seed runs were made, so differences between configurations were assessed with paired participant bootstraps rather than seed-to-seed variation.

### 2.5 Synthetic augmentation and audit

For training only, the 305 real participants were augmented with 1000 synthetic participants generated by a conditional tabular denoising diffusion model (TabDDPM) [38] that modelled the flattened three-visit volume and thickness trajectory, using a cosine noise schedule, v-prediction [39] with min-SNR-5 weighting [40], FiLM conditioning [41] on diagnosis and genotype, and an exponential moving average of weights. We tested whether real and synthetic participants are distinguishable in the trained embedding [42, 43], using the three-visit fold model with the lowest validation loss (fold 4) applied to all 305 real and 1000 synthetic participants, with a classifier two-sample test [44] (five-fold cross-validated logistic regression on the 320-dimensional flattened embedding and on a ten-dimensional raw-feature baseline, 200 label permutations) and a maximum mean discrepancy test [45] (RBF kernel, median-heuristic bandwidth, 300 permutations). Because about 80% of both groups were in the audit model’s training data, the classifier test was repeated separately within its training and its held-out participants. All embedding analyses were restricted to the 305 real participants.

### 2.6 Model variants and validation protocol

#### Held-out-fold protocol

Each real participant was embedded by the single fold-model whose training set excluded them, in evaluation mode with dropout disabled. Month-24 mean atrophy was the negative of the five-region mean of month-24 volumes z-scored against the CN baseline distribution, so that higher values indicate more atrophy.

#### Leakage controls

Feature standardization, group-level edge weights, and the generator were estimated once on the pooled cohort; the generator matters most, because synthetic participants in a training fold carry information about that fold’s held-out real participants. Two retraining controls held architecture, schedule, and seed fixed: a real-only control without synthetic data, and a leak-free control in which a separate generator was trained on each fold’s training participants (244 per fold). Two diagnosis-blind controls were retrained from scratch: a graph-only control that used the pooled graph but retained the diagnostic label in the metadata, and a fully diagnosis-blind control that combined the pooled graph with removal of the diagnostic label from the metadata.

#### Control glossary

*Graph-only* uses pooled diagnostic-group-independent edges while retaining diagnostic metadata; *fully diagnosis-blind* also removes diagnostic metadata; *real-only* excludes synthetic records; *leak-free* denotes a generator fitted within each fold; it does not make every preprocessing step fold-specific.

#### Prospective model

To separate prediction from concurrence, the architecture was retrained on baseline and month-12 visits only, so that month-24 morphometry is never seen by the model. The baseline- to-month-24 cognitive outcome is not strictly subsequent to the inputs, because it overlaps the month-12 imaging visit. A sensitivity analysis therefore used MMSE change from month 12 to month 24, controlling for atrophy and month-12 MMSE, with month-12 scores taken from the ADNI MMSE table and matched by participant identifier. With two visits the encoder retains the velocity term and omits acceleration. The prospective model is diagnosis-blind, with structural-covariance edges recomputed within each training fold, and uses leak-free per-fold augmentation.

#### Site-disjoint validation

ADNI participant identifiers encode the acquisition site (SSS_S_NNNN). The cohort included 47 sites, with a maximum of 19 and a median of 6 participants per site. Because individual sites were too small to provide within-fold neighbourhoods, we assigned whole sites, in descending order of size, to whichever of five folds contained the fewest participants. This yielded 61 held-out and 244 training participants per fold, with no site represented in both sets. The prospective model was retrained on these folds with all other settings unchanged. Site-level inference comprised a sensitivity analysis that omitted each site from the association analysis in turn without retraining, adjustment for 46 site indicators, and a bootstrap that resampled whole sites (2000 draws). The site-disjoint and stratified-split estimates were compared with a paired participant bootstrap of their difference (2000 draws, the same participants resampled for both residuals).

#### Auxiliary-head ablations

The prospective model was retrained with the same folds, generator, graph, optimizer, schedule, and random seeds, either without the tipping head or with the atrophy head alone; the atrophy-head-alone ablation was also repeated within the site-disjoint folds, with the criterion for a retained association (pooled partial *ρ <* 0 with *p <* 0.05) fixed before the retrain. Because every head is downstream of the shared embedding, removing a loss term removes the head’s influence on *h*_node_; we verified that the shared layers received no gradient from the removed loss terms. We compared each ablated residual with the reported one, fit cross-validated ridge regressions predicting the mean month-12 volume-decline rate and mean month-24 volume from the embedding, and repeated the decline analysis with the month-12 volume-decline rate, an input feature, added to the covariates, either as its mean over regions or as all five regional rates.

### 2.7 Off-manifold residual

Fold identity is almost perfectly decodable from the embedding (one-vs-rest AUC *≈* 1.0), so embeddings from independently trained fold-models do not share a coordinate frame. For each held-out participant we therefore identified the ten nearest neighbours among participants embedded by the same fold-model, reconstructed the participant’s embedding as their weighted mean under a self-tuning Gaussian kernel [46] with a seventh-nearest-neighbour bandwidth, and defined the off-manifold residual as the Euclidean reconstruction error divided by the embedding norm; only these scalars were pooled across folds. The residual is related to density-based outlier scores such as the local outlier factor [47]. The prospective, site-disjoint, and ablation models use the un-normalized error, specified before this comparison. Robustness analyses varied the bandwidth (5 to 20 neighbours) and reconstruction count (5 to 30), and examined the un-normalized error, the embedding norm, and adjustment for Mahalanobis distance.

### 2.8 Cognitive endpoints and statistical analysis

#### Cognitive endpoints

Six cognitive measures at baseline and month 24 were matched to the imaging cohort by participant identifier: the Mini-Mental State Examination (MMSE) [48], delayed Rey Auditory Verbal Learning Test recall, category fluency, the Boston Naming Test [49], Trail Making Test Part B [50], and logical-memory delayed recall. This study was not pre-registered and no endpoint hierarchy was specified in advance. The six endpoints were therefore treated as one family and corrected together with the Benjamini–Hochberg procedure, separately for each model and for the pooled and within-AD analyses (Supplementary Table S2); associations with *p <* 0.05 that did not survive this correction are reported as nominal and exploratory. Robustness, ablation, and covariate-adjustment analyses that follow up the MMSE association are reported with unadjusted *p* values and are not confirmatory. Baseline logical memory was taken from the screening visit, at which ADNI-1 administered it. Scores recorded with an ADNI missing-value code at either visit were treated as missing, so the sample size differs by endpoint.

#### Association with decline

We related the residual to 24-month MMSE change with Spearman correlation and with a partial Spearman correlation controlling for month-24 mean atrophy and baseline MMSE, computed by correlating the residuals of rank-on-rank regressions, with percentile bootstrap 95% confidence intervals; an exact precision-matrix estimate was computed for comparison. Analyses were performed pooled and within AD. Robustness analyses replaced mean atrophy with the full set of regional atrophy measures (five regions *×* volume and thickness), recomputed the residual across the neighbourhood grid, and controlled for the input features defining the tipping target.

#### Performance metrics

Discrimination of AD from CN and of AD from all non-AD participants was quantified by the area under the ROC curve (AUC) for the residual, for month-24 mean atrophy, and for both combined in a five-fold cross-validated logistic regression on standardized inputs, with stratified bootstrap 95% confidence intervals (2000 draws); the difference between the combined and atrophy-only AUCs was assessed with a paired participant bootstrap. For MMSE change, nested ordinary-least-squares models with atrophy and baseline MMSE, with and without the residual, gave the incremental *R*^2^ (bootstrap 95% CI) and the residual coefficient’s *p* value, and ten-fold cross-validation gave out-of-sample *R*^2^ and mean absolute error (MAE).

#### Statistical analysis

Group differences were tested with Kruskal–Wallis tests (effect size *ε*^2^ = *H/*(*n −* 1)) and two-sided Mann–Whitney tests, and associations with covariates with Spearman correlation. Multiplicity was controlled with the Benjamini–Hochberg procedure [51] at *q* = 0.05 within the stated analysis families; exploratory regional contrasts used one-sided Mann–Whitney tests and are reported without correction. Permutation nulls used group-stratified resampling [52, 53], and effect sizes (Cohen’s *d*, Cliff’s delta) used bias-corrected bootstrap intervals [54, 55]. Supplementary Table S3 summarizes selected inferential results.

### 2.9 Localization and interpretability

To localize the residual, we recomputed it from each region’s 64-dimensional sub-vector of *h*_node_ and tested AD against the other participants with a one-sided Mann–Whitney test, summarized by the Hodges–Lehmann shift with the confidence interval obtained by inverting the test, in the reported, both diagnosis-blind, and leak-free models. To attribute it to temporal channels, we rebuilt, in the three-visit model (the prospective model stores only its fused embedding), baseline (level), velocity (first differences), acceleration (second difference), and fused representations from the per-visit spatial states and applied the same residual to each, testing group separation and association with MMSE change. For each of the twenty directed edges, attention reconfiguration was the summed absolute change in head-averaged attention across the two visit transitions, correlated with the residual under Benjamini–Hochberg control across edges. For each region pair we computed the Williams–Beer partial information decomposition [56, 57] about month-24 outcome, using Gaussian-copula mutual information [58] and minimum-mutual-information redundancy [59]. Predictive uncertainty and gradient attribution are described in Supplementary Note 6.

### 2.10 Comparison analyses and controls

#### Conventional anomaly scores

Four reference scores were computed on the same input features (five regions *×* seven features *×* three visits) and held-out participants: Mahalanobis distance to the CN distribution with Ledoit–Wolf shrinkage [60], a principal-component reconstruction residual fit on CN participants, the local outlier factor with 20 neighbours [47], and a normative deviation from age- and sex-adjusted CN regressions [61]. Each was evaluated with the partial Spearman analysis described above.

#### Diffusion pseudo-time

As a negative control, the per-visit node-mean embeddings (305 *×* 3 points) were embedded with a diffusion map [62] (self-tuning bandwidth, *α* = 1 normalization), and the per-participant change in the leading non-trivial eigenvector Δ*ψ*_1_ was treated as a diffusion pseudo-time [63] and correlated with month-24 atrophy within AD. We varied the bandwidth (5 to 20 neighbours), recomputed the analysis on the graph-only diagnosis-blind model, and inspected the eigenvalue spectrum and the rotation of the leading eigenspace.

### 2.11 Software and reproducibility

Analyses used Python 3.12.0 with PyTorch 2.3.1, PyTorch Geometric 2.6.1, torch-scatter 2.1.2, torchsparse 0.6.18, NumPy 1.26.4, SciPy 1.14.0, pandas 2.2.2, scikit-learn 1.8.0, statsmodels 0.14.5, matplotlib 3.9.2, and seaborn 0.13.2. All analyses used a master random seed of 42.

## 3 Results

### 3.1 Synthetic augmentation audit and held-out reconstruction

Real and synthetic participants were distinguishable in the trained embedding (classifier AUC 0.66, permutation *p <* 0.005; maximum mean discrepancy *p <* 0.004) but not in the raw features (AUC 0.50, *p* = 0.55; Supplementary Fig. S1). The separability was present both among the audit model’s training participants (AUC 0.65) and among its held-out participants (AUC 0.62, *p* = 0.025; Supplementary Note 2). It therefore arises within the network rather than from a generator failure or from pooling training and held-out participants. The shared embedding reconstructed held-out month-24 volume and thickness with a mean absolute error of 0.14 z-units for both, with similar errors across regions and diagnostic groups; because the three-visit model receives month 24 as input, forecasting was assessed with the prospective model (Section 3.3).

### 3.2 The off-manifold residual separates AD from CN and MCI

The residual differed across diagnostic groups (Kruskal–Wallis *H*(2) = 20.8, *p <* 0.001, *ε*^2^ = 0.068; Fig. 2a,b). Participants with AD had higher residuals (median 0.397) than CN (0.335) and MCI (0.334) participants (both *q <* 0.001), whereas no significant difference was detected between CN and MCI. Higher residuals were nominally associated with more atrophy (Spearman *ρ* = 0.12, 95% CI 0.01 to 0.23, *p* = 0.035), but this association did not survive correction across the four covariates (*q* = 0.14), and the residual was not associated with age, sex, or *APOE ε*4 carrier status (all *q >* 0.4; Fig. 2c). Group separation held across all 25 neighbourhood settings (*H* = 18.0 to 21.7, all *p <* 0.001; Fig. 2d) and in the fully diagnosis-blind retrain (*H* = 21.5, *p <* 0.001), indicating that diagnostic input was not necessary for the separation. Excluding 35 participants with a regional volume change above 15% between consecutive visits reduced but did not remove the separation (*H* = 12.1, *p* = 0.002); the reduction came from 17 participants, 13 with AD, with large and mostly persistent volume losses, whereas excluding those with implausible volume increases left it unchanged. Additional checks on normalization, neighbourhood construction, and augmentation are reported in Supplementary Notes 1 and 2.

**Figure 2.**
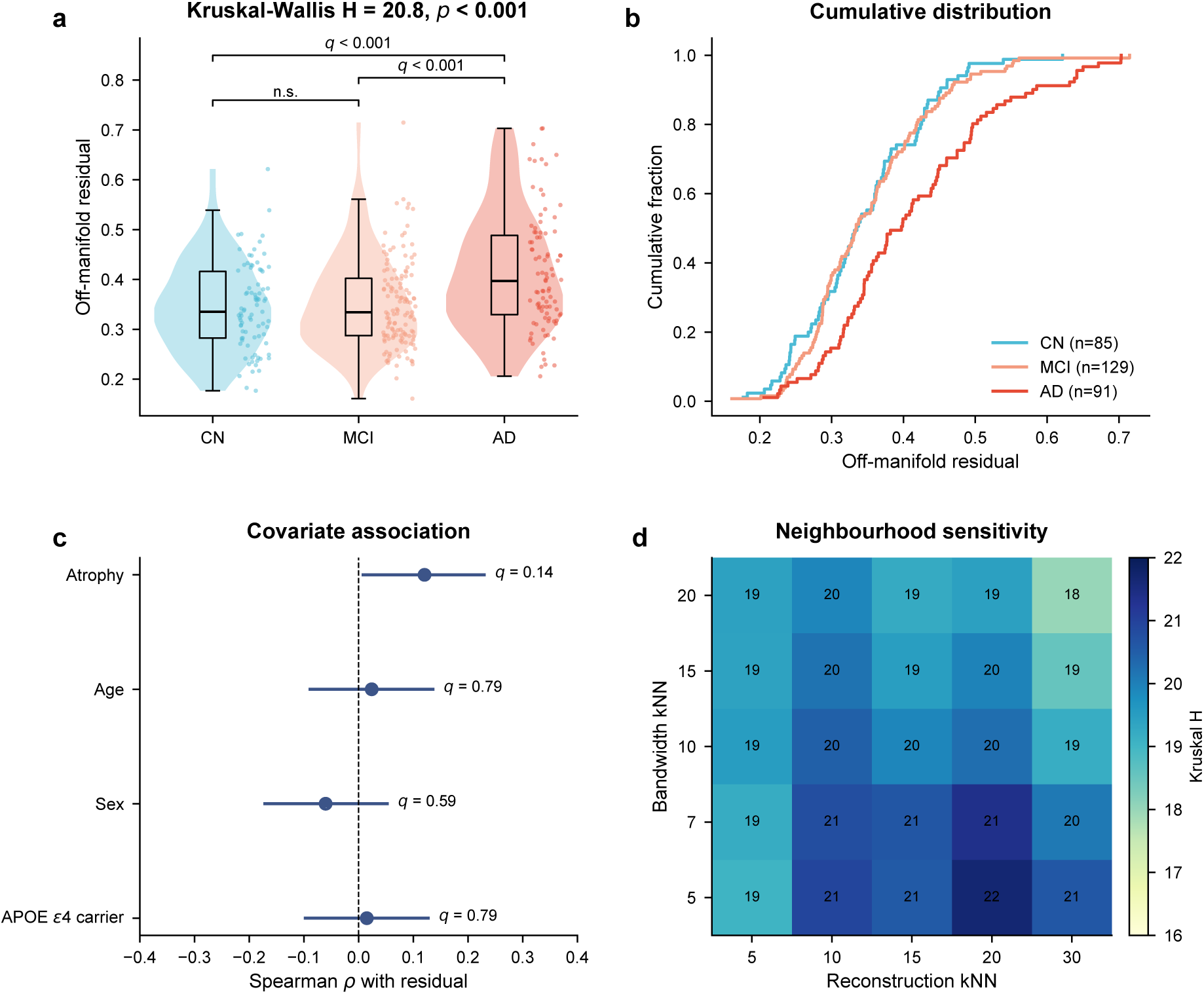
The off-manifold residual separates AD from CN and MCI. **a**, Residual by diagnostic group (violin, box, and individual participants); brackets show two-sided Mann–Whitney tests with Benjamini–Hochberg correction over three contrasts. **b**, Empirical cumulative distributions by group. **c**, Spearman correlation of the residual with atrophy (higher values, more atrophy), age, sex, and *APOE ε*4 carrier status (Fisher 95% CI, uncorrected), with Benjamini–Hochberg-corrected *q* values across the four covariates; the atrophy interval excludes zero (uncorrected *p* = 0.035) although *q* = 0.14. **d**, Kruskal–Wallis *H* across kernel-bandwidth and reconstruction-neighbour settings (rounded; unrounded range 18.0 to 21.7). Residuals are the local reconstruction error divided by the embedding norm; n.s., not significant after Benjamini–Hochberg correction. Three-visit model; *n* = 305.

### 3.3 Association of the residual with cognitive decline

In the three-visit model, higher residuals accompanied steeper 24-month MMSE decline (Spearman *ρ* = *−*0.30, *p <* 0.001; Fig. 3a). Because this model receives the month-24 scan as input, the association is concurrent. It persisted after adjustment for atrophy and baseline MMSE and survived correction across the six endpoints (partial *ρ* = *−*0.25, *p <* 0.001, *q <* 0.001); within AD the estimate was similar but did not survive that correction (partial *ρ* = *−*0.27, 95% CI *−*0.45 to *−*0.07, *p* = 0.011, *q* = 0.067). The pooled association was robust to adjustment for the full set of regional month-24 volume and thickness measures, to the neighbourhood settings, and diagnosis-blind retraining (Supplementary Note 3). Within AD, the estimate remained negative in both augmentation controls but was nominally significant only with the fold-specific generator (*ρ* = *−*0.21, *p* = 0.047) and not with real-only training (*ρ* = *−*0.20, *p* = 0.058; Supplementary Note 2), indicating limited robustness of the within-AD signal across training configurations; these controls do not distinguish a generator effect from sampling variability or from a change in the learned representation.

**Figure 3.**
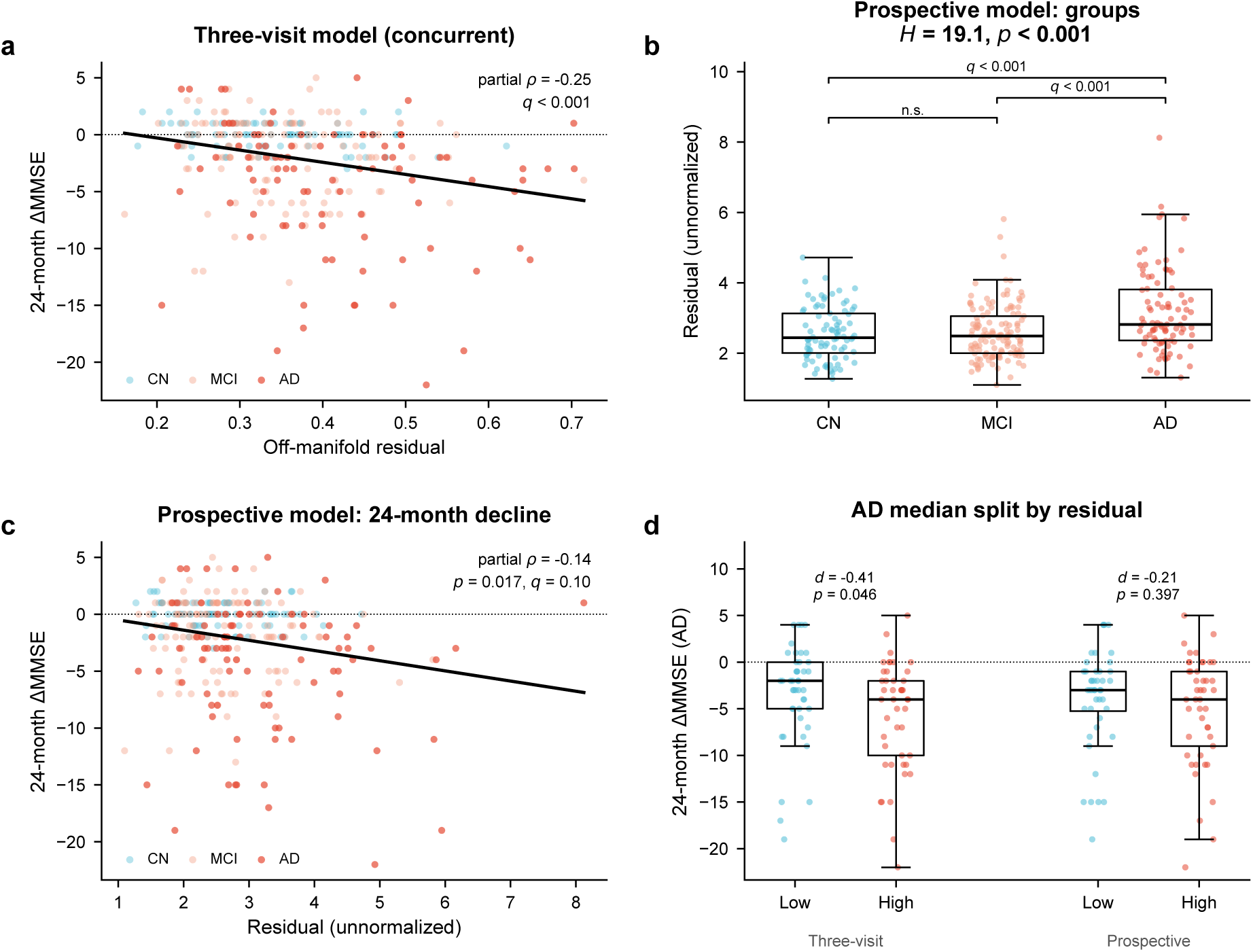
Association of the residual with 24-month cognitive decline. **a**, Three-visit residual against 24-month MMSE change, which is concurrent with the month-24 scan read by this model. **b**, Prospective residual (baseline and month-12 input; un-normalized scale) by diagnostic group; brackets as in Fig. 2. **c**, Prospective residual against 24-month MMSE change; the outcome interval overlaps the month-12 imaging input (for change from month 12 to month 24, see Supplementary Table S1). **d**, MMSE change in the 89 AD participants with month-24 MMSE, split at the median residual, in the three-visit (*p* = 0.046) and prospective (*p* = 0.40) models. Lines in **a** and **c** are descriptive least-squares fits; inference uses partial Spearman correlation controlling for atrophy and baseline MMSE, and *q* in **a** and **c** is Benjamini–Hochberg-corrected across the six cognitive endpoints. Residuals are computed within fold; *n* = 305 (85 CN, 129 MCI, 91 AD), of whom 303 (89 AD) have month-24 MMSE.

The prospective model, which never receives the month-24 scan, reproduced the group separation (*H*(2) = 19.1, *p <* 0.001, *ε*^2^ = 0.063; Fig. 3b) and showed a nominal association with 24-month MMSE change after adjustment for atrophy and baseline MMSE (partial *ρ* = *−*0.14, *p* = 0.017; Fig. 3c) that did not survive correction across the six endpoints (*q* = 0.10) and is therefore exploratory in this split. Because the baseline-to-month-24 change overlaps the month-12 imaging input, we also analysed change from month 12 to month 24 (mean *−*1.2 points), controlling for atrophy and month-12 MMSE. Neither prospective model was associated with this later change (stratified split partial *ρ* = *−*0.05, 95% CI *−*0.16 to 0.07, *p* = 0.42; site-disjoint *ρ* = *−*0.04, *−*0.15 to 0.06, *p* = 0.46; within AD *ρ* = *−*0.16 in both, *p ≥* 0.13; Supplementary Table S1). The prospective evidence therefore concerns change over an interval that includes the imaging window, not decline after it. Without the 35 participants with large visit- to-visit volume changes, the stratified prospective association was not significant (partial *ρ* = *−*0.12, *p* = 0.061), whereas the three-visit and site-disjoint associations were retained (*ρ* = *−*0.22 and *−*0.17; Supplementary Note 1). Retraining with early stopping and checkpoint selection on an inner validation set, so that the held-out fold was not evaluated during training, gave similar estimates (stratified split partial *ρ* = *−*0.15, *p* = 0.009; site-disjoint *ρ* = *−*0.18, *p* = 0.001) and retained the group separation (*H*(2) = 16.1 and 14.2, both *p <* 0.001; Supplementary Note 4). The within-AD association was not significant in this model (*ρ* = *−*0.12, *p* = 0.27), and no endpoint other than MMSE was associated with the residual after correction in any model; in the three-visit model, the Boston Naming Test showed a nominal association that did not survive correction (*ρ* = *−*0.13, *p* = 0.023, *q* = 0.070; Supplementary Table S2). Dividing the prospective residual by the embedding norm increased both group separation and the magnitude of the association with MMSE change (Supplementary Note 4).

Table 3 reports discrimination and decline prediction in conventional metrics. In each model the residual distinguished AD from CN and from all non-AD participants above chance. Adding it to atrophy increased the AUC point estimates by 0.030 to 0.041, but all confidence intervals for these differences included zero. For MMSE change, the residual increased the in-sample *R*^2^ by 0.013 to 0.029 (1.3 to 2.9 percentage points) beyond atrophy and baseline MMSE, and its coefficient had unadjusted *p <* 0.05 in all three models (*p* = 0.027 in the prospective model). The cross-validated mean absolute error was numerically lower with the residual in all three models.

**Table 3.** Discrimination and decline-prediction performance. A. , AUC (95% stratified bootstrap CI) of the residual, month-24 mean atrophy, and both combined (five-fold cross-validated logistic regression); ΔAUC is combined minus atrophy with paired-bootstrap 95% CI. **B**, Prediction of 24-month MMSE change from atrophy and baseline MMSE without and with the residual: in-sample *R*^2^, incremental *R*^2^ (95% bootstrap CI) with the residual coefficient’s *p* value, and ten-fold cross-validated *R*^2^ and mean absolute error (points); arrows separate the models without and with the residual. *n* = 305 for discrimination (AD versus CN, *n* = 176) and *n* = 303 for MMSE change.

| **A** Discrimination (AUC) |  |  |  |  |  |
|---|---|---|---|---|---|
| Model | Contrast | Residual | Atrophy | Residual + atrophy | ΔAUC |
| Three-visit | AD vs CN | 0.67 (0.59, 0.75) | 0.79 (0.72, 0.85) | 0.83 (0.77, 0.89) | 0.041 (−0.001, 0.084) |
|  | AD vs non-AD | 0.66 (0.60, 0.73) | 0.73 (0.67, 0.79) | 0.76 (0.70, 0.82) | 0.035 (−0.004, 0.078) |
| Prospective | AD vs CN | 0.66 (0.58, 0.74) | 0.79 (0.72, 0.86) | 0.83 (0.76, 0.88) | 0.033 (−0.004, 0.074) |
|  | AD vs non-AD | 0.66 (0.59, 0.72) | 0.73 (0.66, 0.78) | 0.76 (0.70, 0.81) | 0.030 (−0.011, 0.071) |
| Site-disjoint | AD vs CN | 0.65 (0.56, 0.73) | 0.79 (0.72, 0.85) | 0.82 (0.76, 0.88) | 0.032 (−0.008, 0.075) |
|  | AD vs non-AD | 0.66 (0.59, 0.72) | 0.73 (0.67, 0.79) | 0.76 (0.70, 0.82) | 0.034 (−0.002, 0.073) |
| **B** 24-month MMSE change |  |  |  |  |  |
| Model | *R*<sup>2</sup> | Δ*R*<sup>2</sup> (95% CI) | *p* | CV *R*<sup>2</sup> | CV MAE |
| Three-visit | 0.231 → 0.255 | 0.024 (0.003, 0.063) | 0.002 | 0.210 → 0.229 | 2.68 → 2.63 |
| Prospective | 0.231 → 0.243 | 0.013 (0.000, 0.049) | 0.027 | 0.210 → 0.213 | 2.68 → 2.66 |
| Site-disjoint | 0.231 → 0.259 | 0.029 (0.004, 0.076) | < 0.001 | 0.210 → 0.229 | 2.68 → 2.63 |

In the three-visit model, AD participants with residuals above the median showed greater MMSE decline than those below it (mean change *−*5.7 versus *−*3.4 points; Cohen’s *d* = *−*0.41, *p* = 0.046), but this split uses the concurrent month-24 scan; in the prospective model the difference was not significant (*d* = *−*0.21, *p* = 0.40; Fig. 3d). These median-split analyses therefore did not support use of the residual for prospective trial enrichment in this cohort (Supplementary Note 7).

### 3.4 The association persists in site-disjoint validation and depends on rate supervision

When the prospective model was retrained on site-disjoint folds, so that no held-out participant shared a site with any training participant, the residual again separated the diagnostic groups (*H*(2) = 19.0, *p <* 0.001, *ε*^2^ = 0.062) and was associated with MMSE change (partial *ρ* = *−*0.20, *p <* 0.001, six-endpoint *q* = 0.003; Fig. 4b), although not significantly within AD (*ρ* = *−*0.21, *p* = 0.053). The association remained significant when each of the 47 sites was omitted in turn (*ρ* from *−*0.23 to *−*0.17; Fig. 4a), after adjustment for 46 site indicators (*ρ* = *−*0.15, *p* = 0.018), and under a site-clustered bootstrap (95% CI *−*0.33 to *−*0.06). The site-disjoint and stratified-split estimates did not differ significantly (paired participant bootstrap difference *−*0.06, 95% CI *−*0.14 to 0.02, *p* = 0.12); because each configuration was trained once with the same seeds, this interval reflects participant sampling, not variability between retrains. Site-level diagnostics are reported in Supplementary Note 4.

**Figure 4.**
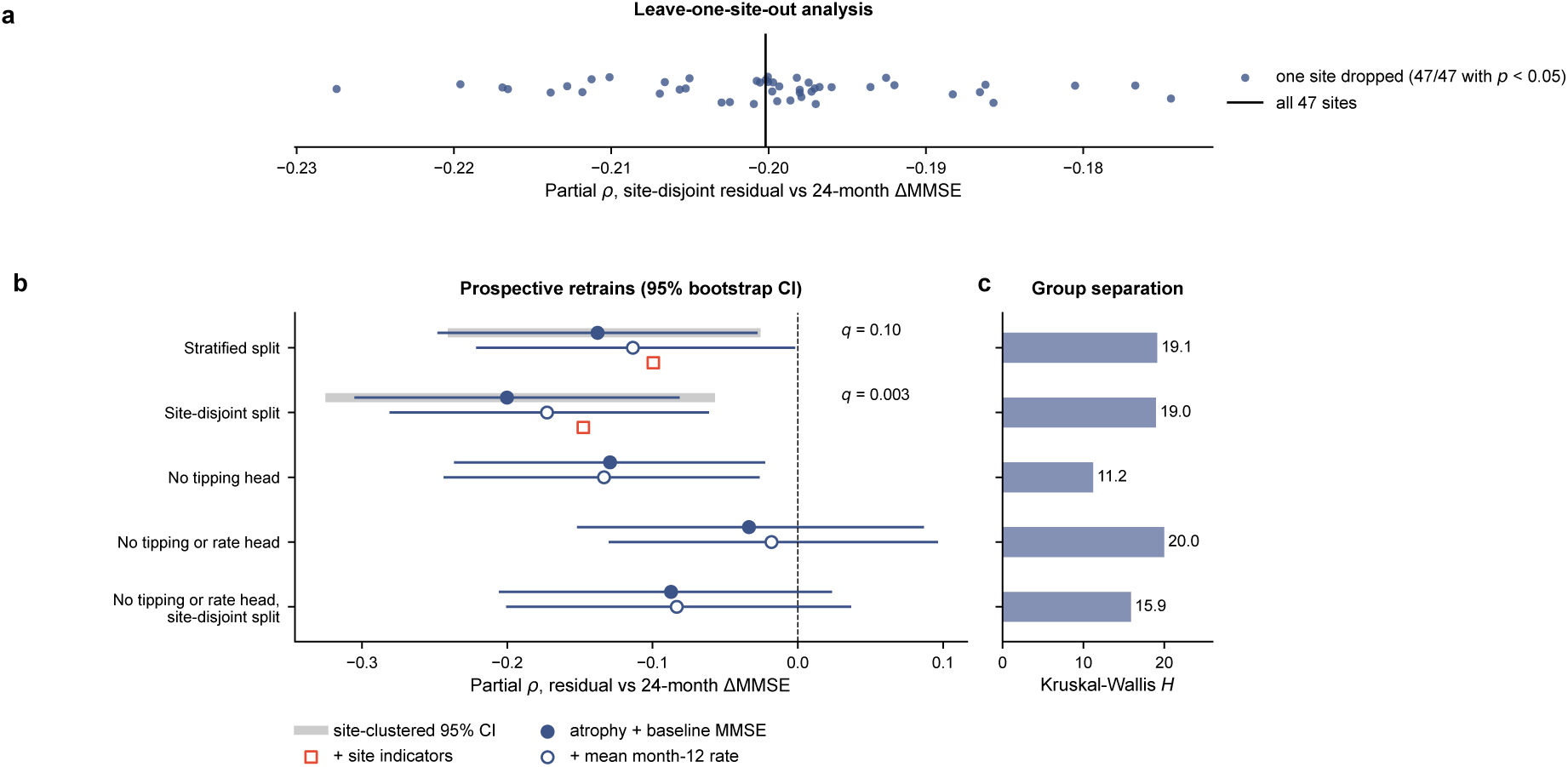
Site-disjoint validation and dependence on rate supervision. **a**, Partial Spearman correlation between the site-disjoint residual and 24-month MMSE change with each of the 47 sites omitted in turn from the association analysis (points; all *p <* 0.05) and with all sites (line); models were not retrained for these omissions. **b**, Partial Spearman correlation for each prospective retrain, controlling for atrophy and baseline MMSE (filled circles; participant bootstrap 95% CI, 2000 draws); the head ablations use the stratified folds unless labelled site-disjoint, and *q* is the Benjamini–Hochberg value across the six cognitive endpoints for the two three-head models; open circles and open squares show separate analyses that also adjust for the mean month-12 volume-decline rate or for 46 site indicators, respectively. Grey bars show site-clustered bootstrap 95% CIs for the stratified-split and site-disjoint models. **c**, Kruskal–Wallis *H* for group separation in each retrain. *n* = 303 for **a**,**b** and *n* = 305 for **c**.

The ablations showed which training objective this association depends on (Fig. 4b,c). Without the tipping head, the residual was strongly correlated with that of the reported model (Spearman *ρ* = 0.91) and remained associated with decline (partial *ρ* = *−*0.13, *p* = 0.025); group separation was weaker (*H*(2) = 11.2, *p* = 0.004, *ε*^2^ = 0.037) but retained the AD-specific pattern. Because the tipping target is a near-deterministic function of two input features, this effect most likely reflects regularization of the shared representation, not information about a clinical tipping point. Removing the rate-supervision head as well preserved group separation (*H*(2) = 20.0), but the association with decline was reduced to near zero and was no longer significant (*ρ* = *−*0.03, *p* = 0.56). The same ablation within the site-disjoint folds gave the same outcome (Table 4): group separation was preserved (*H*(2) = 15.9, *p <* 0.001, *ε*^2^ = 0.052), but the association with MMSE change was no longer significant (*ρ* = *−*0.09, *p* = 0.13) and was weaker than in the three-head site-disjoint model (paired-bootstrap difference *−*0.11, 95% CI *−*0.22 to *−*0.01). The site-disjoint association also depends on rate supervision. Consistent with this, the cross-validated decodability of the month-12 volume-decline rate from the embedding fell from *R*^2^ = 0.92 to 0.68, whereas month-24 volume showed similar decodability (*R*^2^ = 0.83 and 0.84). Because the rate target derives from an input feature, which is itself associated with decline (partial *ρ* = *−*0.17, *p* = 0.003), we added the mean month-12 rate to the covariates. After this adjustment, the associations with MMSE change remained significant for the site-disjoint (*ρ* = *−*0.17, *p* = 0.003) and no-tipping (*ρ* = *−*0.13, *p* = 0.021) models, whereas the stratified-split association was attenuated to the margin of significance (*ρ* = *−*0.11, *p* = 0.049). Adjusting instead for all five regional month-12 rates gave the same pattern (site-disjoint *ρ* = *−*0.18, *p* = 0.002; no-tipping *ρ* = *−*0.14, *p* = 0.016; stratified *ρ* = *−*0.12, *p* = 0.036; Supplementary Note 5).

**Table 4.** Rate-supervision ablation within stratified and site-disjoint folds. Partial Spearman correlation between the residual and 24-month MMSE change, controlling for atrophy and baseline MMSE (participant bootstrap 95% CI, 2000 draws, as in Fig. 4b; *n* = 303 pooled, *n* = 89 AD), and Kruskal–Wallis *H* for group separation (*n* = 305). In the atrophy-loss-only models neither the rate-supervision head nor the tipping head contributes to the objective.

| Folds | Training objective | Pooled partial *ρ* (95% CI) | *p* | Within-AD *ρ* (*p*) | *H*(2) |
|---|---|---|---|---|---|
| Site-disjoint | three heads | −0.20 (−0.30, −0.08) | < 0.001 | −0.21 (0.053) | 19.0 |
| Site-disjoint | atrophy loss only | −0.09 (−0.21, 0.02) | 0.13 | −0.11 (0.31) | 15.9 |
| Stratified | atrophy loss only | −0.03 (−0.15, 0.09) | 0.56 | 0.02 (0.87) | 20.0 |

### 3.5 Comparison with conventional anomaly scores

Among four conventional anomaly scores computed on the same input features, the normative deviation, local outlier factor, and Mahalanobis distance were also associated with pooled concurrent decline beyond atrophy and baseline MMSE (partial *ρ* = *−*0.24 to *−*0.21, all *p <* 0.001), and the principal-component residual was not (Supplementary Fig. S2). Within AD, the off-manifold residual (*ρ* = *−*0.27, *p* = 0.011) and the normative deviation (*ρ* = *−*0.24, *p* = 0.027) reached nominal significance, whereas the local outlier factor, Mahalanobis distance, and principal-component residual did not (*|ρ| ≤* 0.14, *p ≥* 0.19); the residual performed comparably to the best conventional score, and these results do not establish a difference between the scores.

### 3.6 The atypicality is distributed across salience and executive-control regions

When the residual was recomputed separately for each region, the dorsal anterior insula (dAI), the hub of the salience network, showed the largest AD-versus-rest shift in the reported model (one-sided Mann–Whitney *p* = 0.005, uncorrected; Fig. 5a). In the graph-only diagnosis-blind model all five regions differed nominally, with the smallest *p* value for the dAI (*p <* 0.001; dlPFC and SPL, *p* = 0.003), and in the fully diagnosis-blind model the dAI difference had *p* = 0.008. Under leak-free augmentation, the dAI (*p* = 0.039), vlPFC (*p* = 0.035), and dACC (*p* = 0.026) showed nominal differences (Supplementary Note 2). These exploratory results indicate a pattern distributed across salience and executive-control regions rather than a stable ranking of a single region.

**Figure 5.**
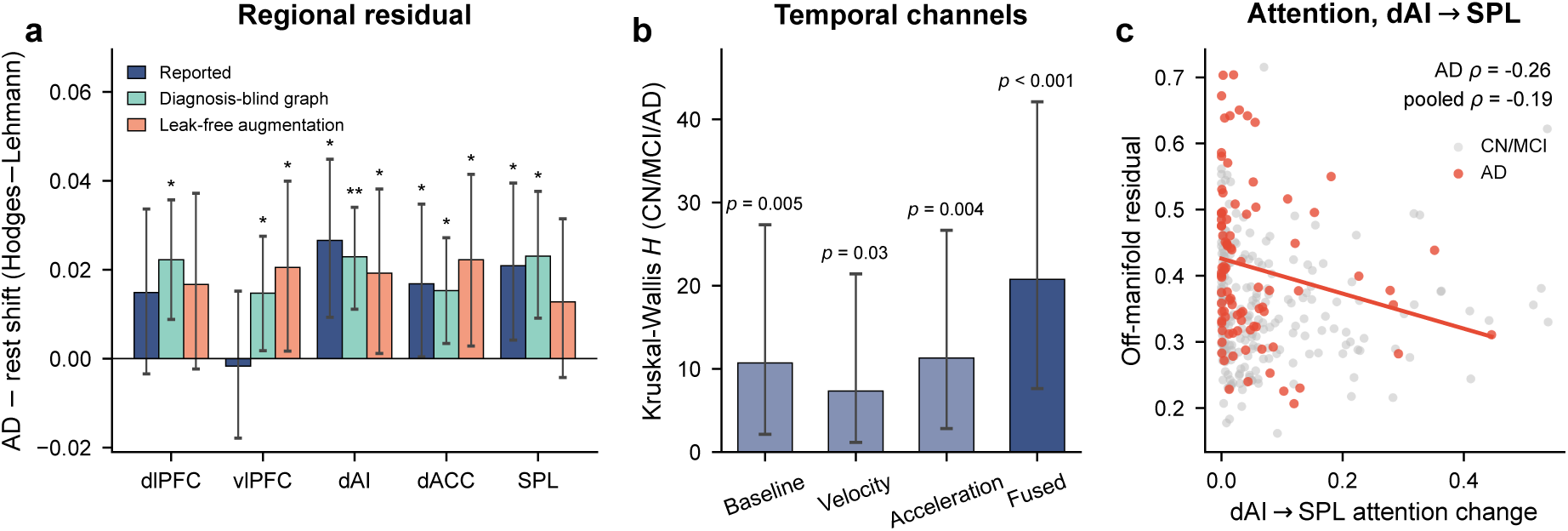
Localization of the residual in the three-visit model. **a**, Per-region residual, AD minus other participants, in the reported, graph-only diagnosis-blind (pooled edges, diagnostic metadata retained), and leak-free-augmentation models: Hodges–Lehmann shift with the 90% confidence interval obtained by inverting the Mann–Whitney test, whose lower bound is the one-sided 95% bound matching the test; asterisks, one-sided Mann–Whitney *p <* 0.05 (single) and *p <* 0.001 (double), uncorrected, and an asterisk appears only where the lower bound exceeds zero. **b**, Group separation (Kruskal–Wallis *H*, bootstrap 95% CI) of the residual computed from each temporal channel and from the fused embedding. **c**, dAI*→*SPL attention reconfiguration against the residual; the line is a least-squares fit within AD. Residuals are computed within fold; *n* = 305.

Each temporal channel separated the groups on its own (baseline *H* = 10.7; velocity *H* = 7.3, *p* = 0.026; acceleration *H* = 11.3), and the fused embedding separated them most strongly (Fig. 5b). Velocity showed the weakest group separation but was the only single channel with a nominally significant association with MMSE change (partial *ρ* = *−*0.13, *p* = 0.026), consistent with a contribution of first-order change to the two-visit prospective model.

Higher residuals were associated with smaller changes in attention on the dAI*→*SPL edge (Spearman *ρ* = *−*0.19, *q* = 0.005; within AD *ρ* = *−*0.26, *p* = 0.014; Fig. 5c), and six of the eight directed edges incident to the dAI remained significant after FDR correction. These findings describe learned attention over structural-covariance edges and should not be interpreted as altered physiological connectivity. A partial information decomposition identified the dAI–SPL pair as carrying the largest synergistic information about month-24 atrophy (0.220 nats; Supplementary Fig. S3). Monte Carlo dropout uncertainty was positively correlated with the residual, whereas Integrated Gradients attribution was approximately uniform across regions (Supplementary Note 6).

### 3.7 A diffusion pseudo-time readout is a graph artifact

At the default bandwidth, a diffusion pseudo-time computed from the same embedding was associated with atrophy within AD (Spearman *ρ* = *−*0.30, *p* = 0.004) but failed the controls that the residual passed (Supplementary Fig. S4). Its sign reversed across the 5-to-20-neighbour bandwidth range, it was not significant after graph-only diagnosis-blind retraining (*ρ* = *−*0.09, *p* = 0.39), and at the default bandwidth (*k* = 15) the leading diffusion eigenvalues were near-degenerate (smallest leading eigengap 0.006), so that the leading two-dimensional eigenspace rotated by up to 86*^◦^* across bandwidths. The pseudo-time is therefore a graph artifact and was not used as a biomarker.

## 4 Discussion

A geometric measure of trajectory atypicality, read from a learned spatio-temporal embedding of morphometry in executive-control and salience-network regions, distinguished AD from CN and MCI and was only nominally correlated with atrophy, an association that did not survive correction. Residuals from the two-visit model, which does not receive the month-24 scan, were associated with MMSE change from baseline to month 24 when whole acquisition sites were withheld from training; with the stratified split the association was nominal and did not survive correction across the six cognitive endpoints. Neither split was associated with change from month 12 to month 24, the interval that follows all imaging inputs, so the evidence concerns cognitive change during the imaging window rather than decline after it. MMSE was not used as a training target.

The residual characterizes trajectory atypicality relative to the learned embedding geometry, not the amount of tissue loss alone. Its principal adjusted associations with cognitive change were modest (partial *ρ* approximately *−*0.14 to *−*0.25). Adding the residual to models containing the atrophy measure produced small numerical improvements in discrimination, but all ΔAUC confidence intervals included zero; the incremental explained variance in cognitive change was small (1.3 to 2.9 percentage points). A geometric scoring rule does not imply an unsupervised representation: the off-manifold residual is computed geometrically, but the embedding in which distances are measured is shaped by the training targets, including explicit rate supervision. The ablations therefore support interpreting the residual as atypicality within a rate-supervised representation, not as a measure of disease dynamics that is independent of the training objective. Its association with cognitive change may partly reflect rate information carried by that representation. Adjustment for the mean month-12 rate, or for all five regional rates, preserved the association in the site-disjoint and no-tipping configurations, but left the stratified prospective association only at the margin of significance (*p* = 0.049); this does not establish that the residual captures a biological process independent of rate.

The localization analyses give the marker an anatomical interpretation. The dorsal anterior insula is a hub of the salience network [22], whose connectivity changes with AD severity, increasing at the very mild stage and declining thereafter [20, 21], and the regional residual, attention reconfiguration, and information synergy all point to the insula–parietal connection. Because these readouts derive from the same embedding and outcome, their agreement indicates internal consistency rather than independent replication, and the leakage controls place the effect in the salience and executive-control system as a whole, not in the insula alone. The localization comes from the three-visit model and the evidence on MMSE change without the month-24 scan from the two-visit model, which carries level and rate but not acceleration; the channel decomposition links the two through the velocity channel.

The pseudo-time result is also a methodological finding. A coordinate that resembled a progression axis under default settings was a graph artifact, and the bandwidth and diagnosis-blind controls that exposed it can be applied to any embedding-derived progression measure.

Among prior brain-network models, BrainNetCNN [13] and BrainGNN [14] learn from connectivity without temporal derivatives, and the Brain Network Transformer [15] extends attention-based brain modelling at a single timepoint. Longitudinal approaches such as TADPOLE entries [19], SuStaIn [18], and event-based progression scores [17] encode time but flatten the graph or rely on hand-crafted features. The model used here combines graph attention with temporal differences, and the off-manifold residual, which is related to density-based outlier scores [47], cannot be computed by a single-timepoint model.

This study has limitations. First, synthetic participants are distinguishable from real participants in the trained embedding, and the generator of the three-visit model was fit without a fold split; the nominal within-AD significance and the regional ranking varied across augmentation configurations, whereas the prospective and site-disjoint models use per-fold generators. Embedding-level audits of this kind should accompany generative augmentation of medical data [43, 64]. Second, the reported models include auxiliary heads whose targets derive from the inputs. The tipping head was not necessary for the association with MMSE change, although its removal weakened diagnostic group separation, whereas removing rate supervision as well left no significant association in either the stratified or the site-disjoint folds, which ties the marker to explicit rate supervision. The two-head configuration evaluated here requires validation, and supervision defined over a subsequent, non-overlapping interval should be examined. Third, all data come from ADNI. Site-disjoint validation tests transfer across scanners and recruitment streams, but ADNI sites share acquisition protocols, inclusion criteria, and quality control, so replication in an independent cohort such as AIBL [65] or the UK Biobank [66, 67] is required; the structural-covariance edges also assume linear inter-regional covariance, and multi-site harmonization remains a concern [68]. Fourth, the cognitive evidence rests on a single instrument with modest effect sizes, and no endpoint hierarchy was pre-registered: only MMSE change was associated with the residual after correction across six endpoints, and not in the stratified prospective split; within-AD prognosis and trial enrichment were not established, and *APOE ε*4 carrier contrasts in prediction residuals varied in direction across regions and diagnostic groups, with none significant after correction for 15 comparisons (Supplementary Fig. S5) [8, 25]. Fifth, the residual was nominally associated with atrophy, and the absence of a corrected association does not establish independence or equivalence; the adjustment set did not include education, intracranial volume, amyloid or tau status, or white-matter hyperintensity burden; and the model excludes medial temporal regions, so we make no claim of incremental value over hippocampal atrophy.

A falsifiable prediction is that the residual contributes information about subsequent cognitive decline beyond the month-12 rate in an independent cohort. This should be tested with a prespecified scoring pipeline, covariates available at the prediction time, and an outcome interval beginning after the imaging-input window. If, in an adequately powered external evaluation, adjustment for the month-12 rate reduces the residual–decline association to near zero, with a confidence interval excluding a prespecified clinically meaningful effect, and the residual provides no incremental out-of-sample predictive value, the claim that the geometric score adds prognostic information beyond rate would be undermined. A nonsignificant result alone would not falsify the claim, and such a finding would count against the added prognostic value of the residual, not against its definition.

Several extensions follow directly. Characterizing participants with high residuals by genetic, fluid, and amyloid profiles would connect the geometric marker to the precision-medicine [69] and biological-staging [3] frameworks. Extending the temporal encoder to irregular visit spacing would allow the residual to be computed from routine clinical follow-up, and functional imaging of the insula–parietal connection would test the localization in an independent modality. Validation of the two-head model, a baseline-only variant, and an independent cohort are the most direct next tests of the marker.

## 5 Conclusion

We derived an off-manifold residual from a spatio-temporal graph attention network trained on longitudinal structural MRI; the residual measures how atypical a participant’s morphometric trajectory is. The residual was elevated in AD and, when the network was trained with rate supervision, showed a modest association with baseline-to-month-24 MMSE change that survived correction across six cognitive endpoints in the three-visit and site-disjoint models but not in the stratified prospective model. The association did not extend to change from month 12 to month 24, after the last imaging input. These findings support further evaluation of morphometric trajectory atypicality as a candidate marker; independent validation with an outcome window after the imaging inputs is needed to establish prognostic utility.

## Supporting information

TRIPOD+AI reporting checklist

## Ethics statement

This study is a secondary analysis of de-identified data from the Alzheimer’s Disease Neuroimaging Initiative. ADNI was approved by the institutional review boards of all participating sites, and written informed consent was obtained from every participant or their authorized representative at enrolment. No new data were collected for this work.

## Data and code availability

Imaging, genetic, and cognitive data are available from the ADNI database (https://adni.loni.usc. edu) to researchers who accept the ADNI data-use agreement. The analysis and figure code, together with the summary-level source-data tables, are available from the corresponding author upon request. In accordance with the ADNI Data Use Agreement, no participant-level raw or derived data are re-distributed by the authors; participant-level derived data (per-subject residuals and embeddings) and trained model weights will be offered to ADNI through its derived-data submission process, and can be regenerated from ADNI data with the analysis code.

## Author contributions

**Arman Dinarvand:** Conceptualization, Methodology, Software, Validation, Formal analysis, Investigation, Resources, Data curation, Writing – original draft, Visualization, Project administration. **Amirali Moradniaei:** Methodology, Software, Validation, Formal analysis, Data curation, Writing – original draft, Visualization. **Elaheh Amini:** Supervision, Writing – review & editing, Validation. **Keivan Borna:** Supervision, Writing – review & editing, Validation.

## Acknowledgements

Data collection and sharing for the Alzheimer’s Disease Neuroimaging Initiative (ADNI) is funded by the National Institute on Aging (National Institutes of Health Grant U19 AG024904). The grantee organization is the Northern California Institute for Research and Education. In the past, ADNI has also received funding from the National Institute of Biomedical Imaging and Bioengineering, the Canadian Institutes of Health Research, and private sector contributions through the Foundation for the National Institutes of Health (FNIH) including generous contributions from the following: AbbVie, Alzheimer’s Association; Alzheimer’s Drug Discovery Foundation; Araclon Biotech; BioClinica, Inc.; Biogen; Bristol-Myers Squibb Company; CereSpir, Inc.; Cogstate; Eisai Inc.; Elan Pharmaceuticals, Inc.; Eli Lilly and Company; EuroImmun; F. Hoffmann-La Roche Ltd and its affiliated company Genentech, Inc.; Fujirebio; GE Healthcare; IXICO Ltd.; Janssen Alzheimer Immunotherapy Research & Development, LLC.; Johnson & Johnson Pharmaceutical Research & Development LLC.; Lumosity; Lundbeck; Merck & Co., Inc.; Meso Scale Diagnostics, LLC.; NeuroRx Research; Neurotrack Technologies; Novartis Pharmaceuticals Corporation; Pfizer Inc.; Piramal Imaging; Servier; Takeda Pharmaceutical Company; and Transition Therapeutics.

---

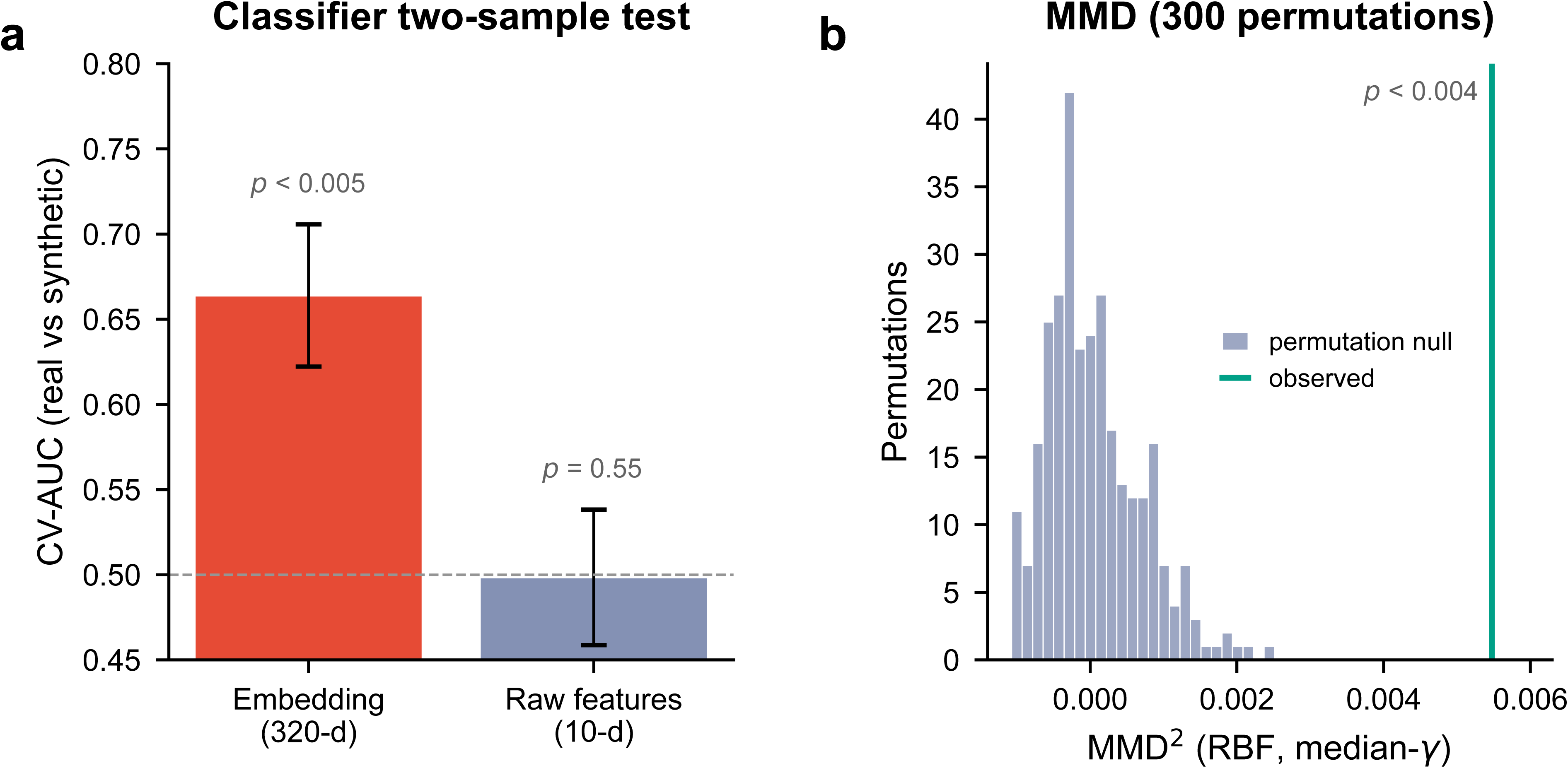

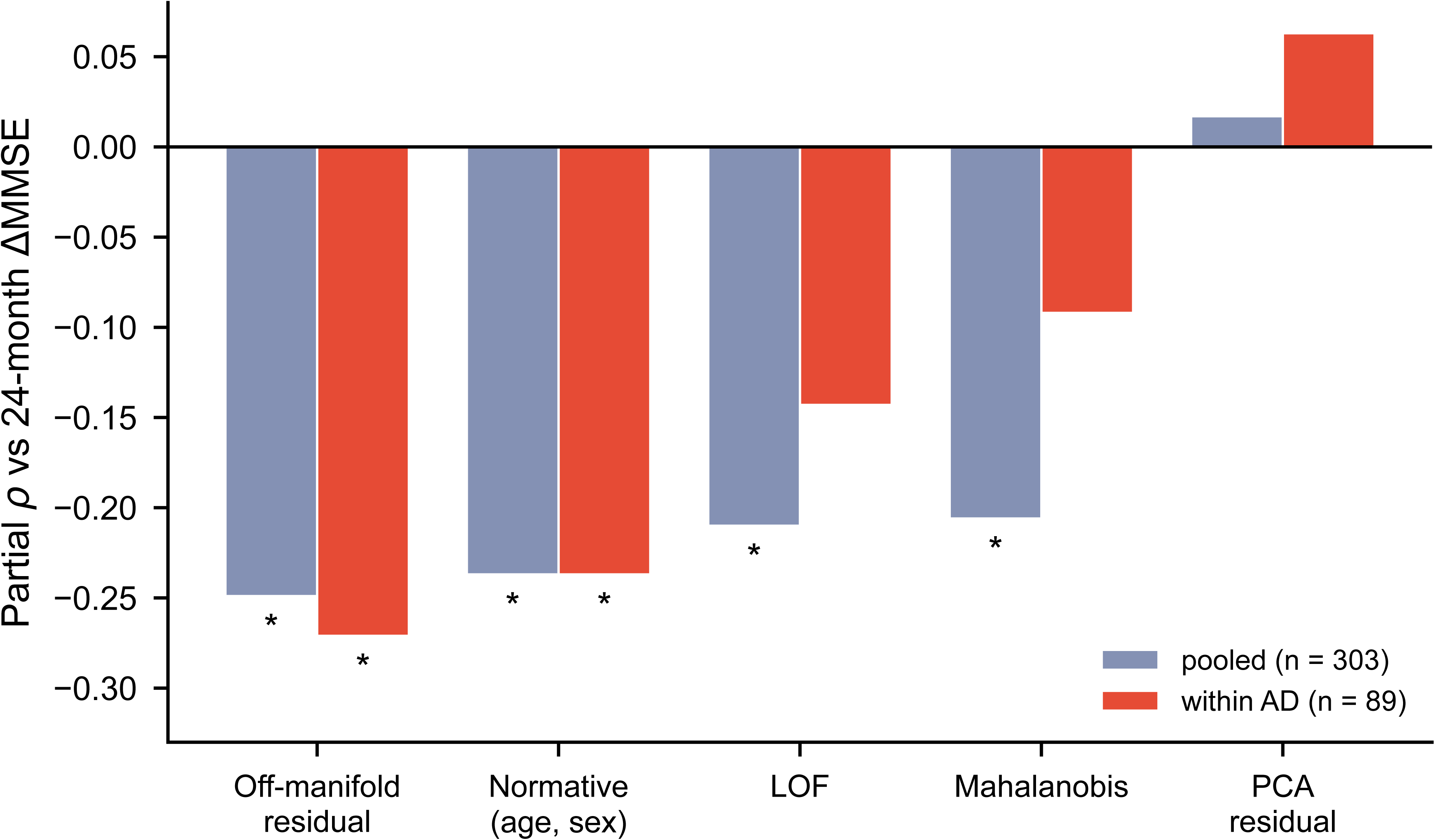

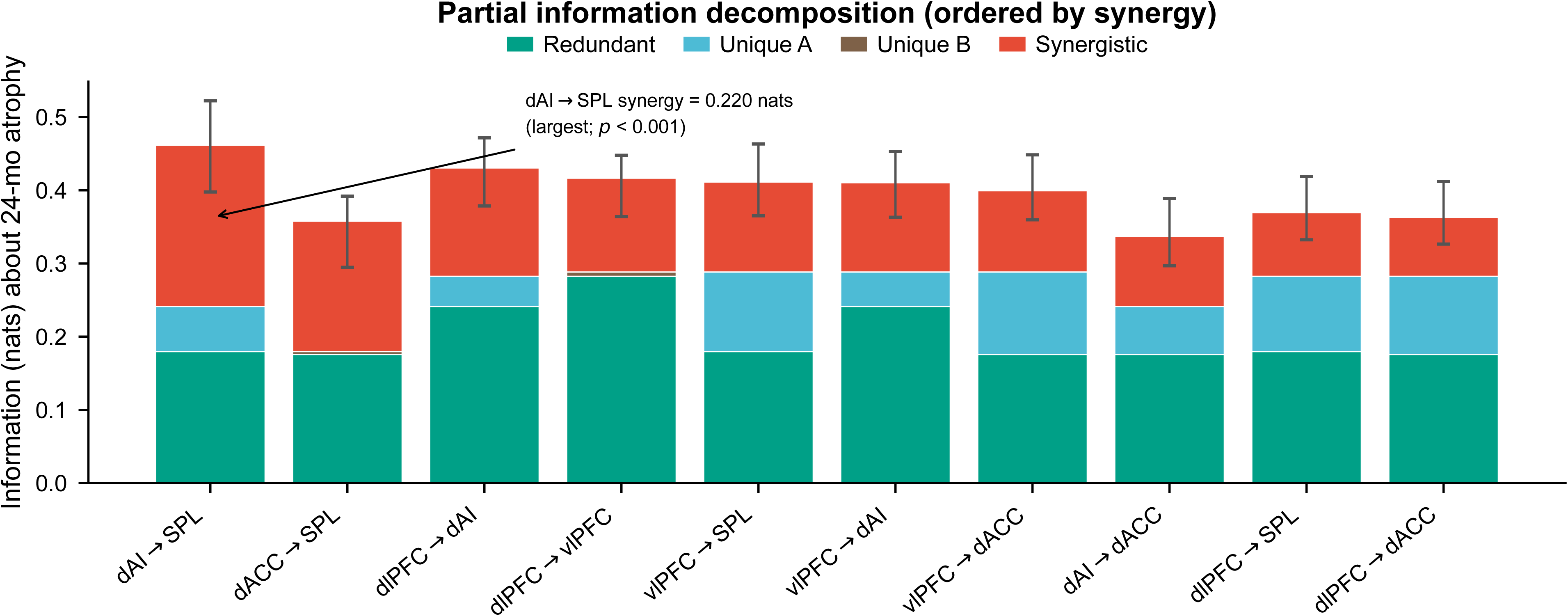

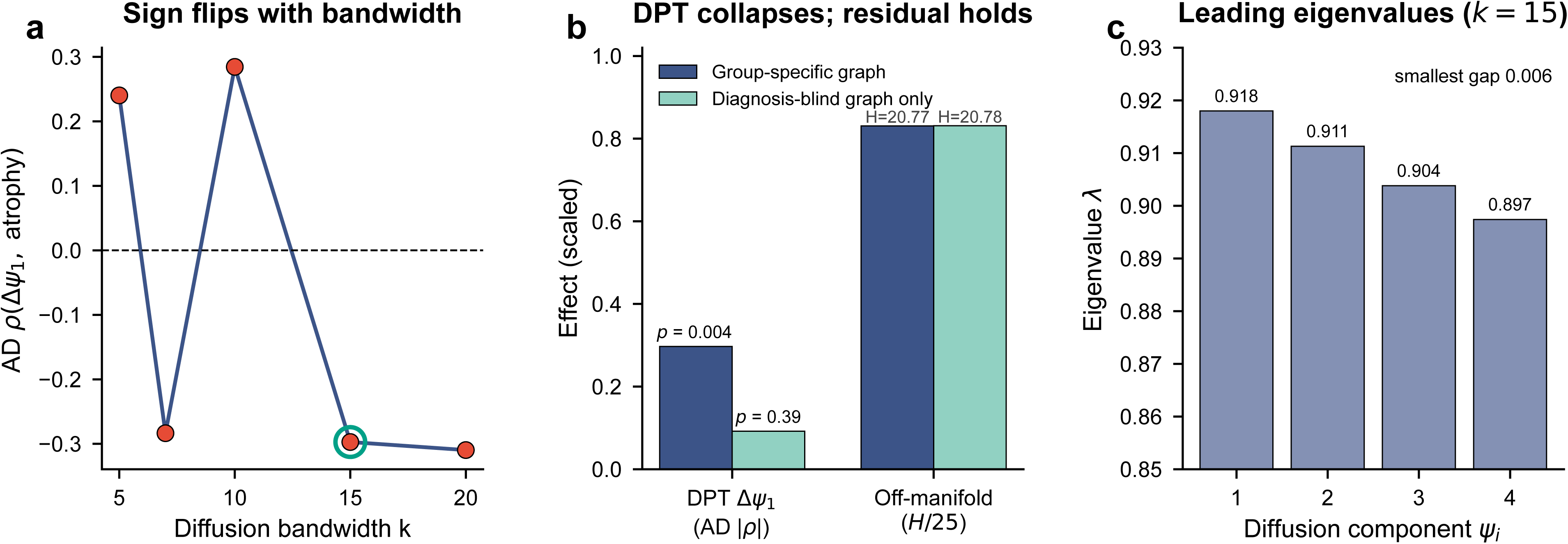

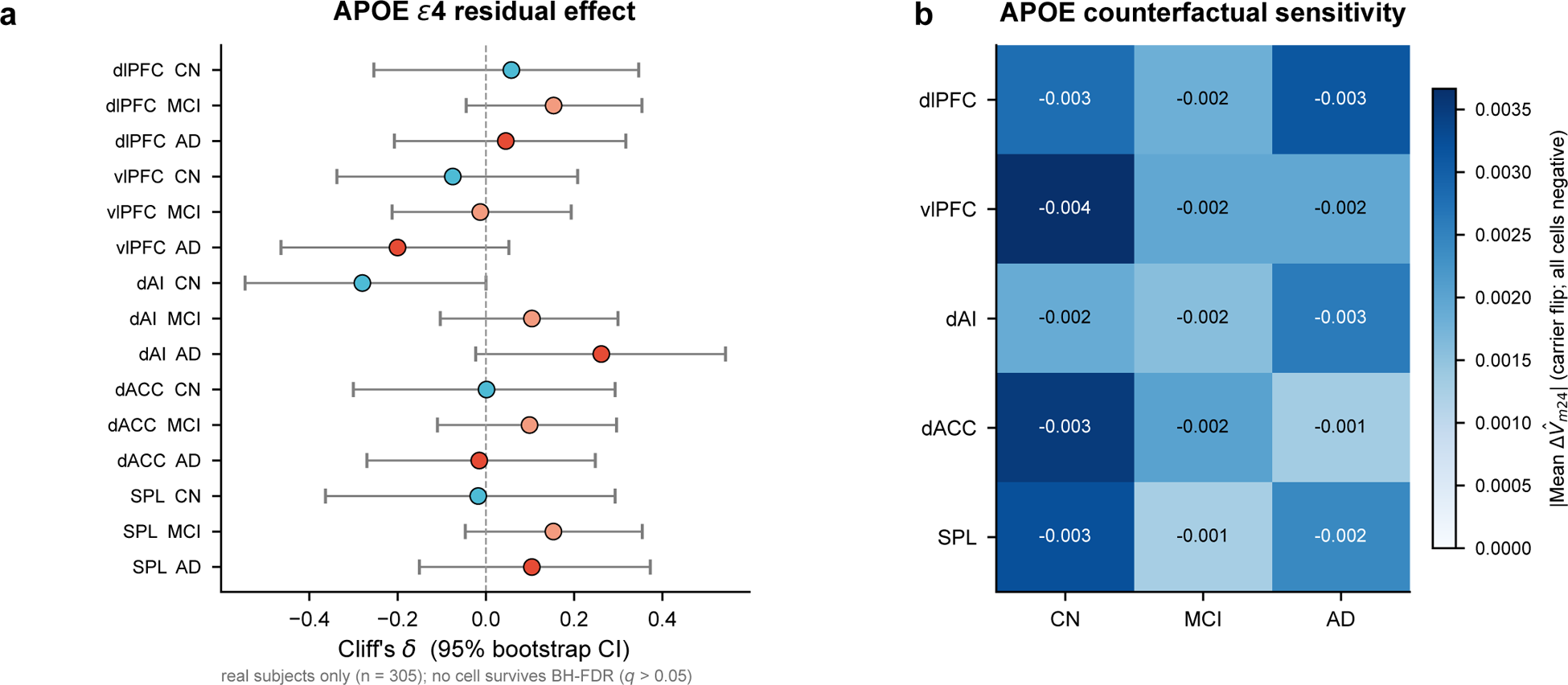

