## Supplementary Files for "A Candidate Trajectory-Atypicality Marker of Concurrent Cognitive Change in Alzheimer’s Disease from a Spatio-Temporal Graph Attention Network"

This file contains Supplementary Notes 1–7, Supplementary Figures S1–S5, and Supplementary Tables S1–S2. Section, figure, and table numbers without an S prefix refer to the main text.

#### Supplementary Note 1. Robustness of the group separation

The un-normalized reconstruction error also separated the diagnostic groups ($H = 17.2$, $q = 0.001$), whereas the embedding norm did not differ significantly by group ($H = 1.8$, $p = 0.40$), indicating that normalization alone does not account for the separation. Each participant's ten nearest neighbours belonged to the same fold as that participant (0 of 305 had a cross-fold neighbour), so restricting neighbourhoods to the fold left the statistic unchanged ($H = 20.767$). In the fully diagnosis-blind retrain, the median residual was 0.399 in AD versus 0.348 in CN and 0.344 in MCI. In the graph-only diagnosis-blind retrain, the group statistic was $H = 20.78$ (AD median 0.328 versus 0.249 in CN and 0.281 in MCI), which coincides to one decimal place with that of the reported model ($H = 20.77$) although the two models were trained separately. Adjusting the residual for Mahalanobis distance to the CN distribution left the group separation intact ($H = 18.1$, $p < 0.001$). The within-fold residual was concordant with the residual of the leak-free model (Spearman $\rho = 0.81$ pooled, 0.85 within AD).

Thirty-five participants had at least one regional volume change between consecutive visits exceeding 15% (65 region-intervals). In 18 of them (8 CN, 8 MCI, 2 AD) at least one change was an increase, which is implausible over one year and suggests segmentation error; the other 17 (13 AD, 3 MCI, 1 CN) had decreases only, most of which persisted at the next visit. Excluding all 35 reduced group separation in the three-visit model from $H = 20.8$ to $H = 12.1$ ($p = 0.002$), and similarly in the prospective ($H = 12.3$) and site-disjoint ($H = 12.0$) models. Excluding only the 18 participants with increases left the separation unchanged ($H = 20.9$); the 17 participants with large losses had residuals at the 95th percentile and accounted for the reduction. Without the 35 participants, the association with 24-month MMSE change was retained in the three-visit (partial $\rho = -0.22$, $p < 0.001$) and site-disjoint ($\rho = -0.17$, $p = 0.006$) models but not in the stratified prospective model ($\rho = -0.12$, $p = 0.061$); within AD, no association was significant (three-visit $\rho = -0.20$, $p = 0.092$), and the month-12-to-month-24 association remained null. Recomputing the residual without these participants as neighbours gave the same pattern.

#### Supplementary Note 2. Augmentation controls

In the real-only control, the residual separated the groups ($H = 18.4$, $p < 0.001$) and was associated with pooled MMSE change after adjustment for atrophy and baseline MMSE (partial $\rho = -0.17$, $p = 0.004$); the within-AD association was $\rho = -0.20$ ($p = 0.058$). Under leak-free per-fold augmentation, the corresponding values were $H = 21.8$ ($p < 0.001$; AD median 0.403 versus 0.347 in CN and 0.355 in MCI), pooled $\rho = -0.21$ ($p < 0.001$), and within-AD $\rho = -0.21$ ($p = 0.047$). In the regional analysis under leak-free augmentation, the dAI ($p = 0.039$), vlPFC ($p = 0.035$), and dACC ($p = 0.026$) showed nominally significant differences (one-sided, uncorrected $p$ from 0.026 to 0.039). Group separation and the pooled association with MMSE change persisted in both augmentation controls; within AD, the estimates remained negative but were significant only under leak-free augmentation, and the regional ranking varied across augmentation configurations.

The synthetic-data audit (Supplementary Fig. S1) used the three-visit fold-4 model, whose training data contained about 80% of both the real and the synthetic participants. Repeated separately within that model's training participants (243 real, 801 synthetic) and within its held-out participants (62 real, 199 synthetic), the classifier gave AUC 0.65 ($p < 0.005$) and 0.62 ($p = 0.025$), respectively. The separability is therefore present in participants the audit model was not trained on and is not explained by pooling training and held-out participants.

#### Supplementary Note 3. Robustness of the concurrent decline association

Replacing mean atrophy with the full set of regional atrophy measures (five regions × month-24 volume and thickness) gave a pooled partial $\rho = -0.27$ ($p < 0.001$) and a within-AD $\rho = -0.32$ ($p = 0.004$). Across a $5 \times 4$ grid of bandwidth and reconstruction neighbours, the pooled partial $\rho$ ranged from $-0.25$ to $-0.22$ (all $p < 0.05$); the within-AD $\rho$ was negative and significant at every setting. In the fully diagnosis-blind model the pooled partial $\rho$ was $-0.25$ ($p < 0.001$) and the within-AD $\rho$ was $-0.28$ ($p = 0.008$). The exact precision-matrix estimate reproduced the rank-residual estimate ($\rho = -0.249$). The ten input features that define the tipping target (five regions × month-12 volume and within-subject volume slope) accounted for 14% of the residual's variance pooled and 29% within AD (linear models); after adjustment for them and baseline MMSE, the pooled association remained $\rho \approx -0.25$ ($p < 0.001$; incremental $R^2 = 0.024$, $p = 0.002$), with a within-AD $\rho = -0.20$ ($p = 0.08$). With 46 site indicators added, the pooled three-visit association was $\rho = -0.21$ ($p < 0.001$).

#### Supplementary Note 4. Prospective and site-disjoint models

Dividing the reconstruction error by the embedding norm gave $H = 27.2$ and partial $\rho = -0.16$ ($p = 0.005$) in the prospective model, and $H = 23.8$ and $\rho = -0.16$ ($p = 0.007$) in the site-disjoint model; the main text reports the un-normalized residual, specified before these comparisons. Site-disjoint folds contained 15 to 24 AD participants each. Within AD, the site-disjoint association was $\rho = -0.21$ ($p = 0.053$); within-AD results were not consistently significant across model configurations. In the stratified-split prospective model, adding 46 site indicators attenuated the association to $\rho = -0.10$ ($p = 0.11$), whereas its site-clustered bootstrap interval excluded zero (95% CI $-0.24$ to $-0.03$). Across the 17 sites with at least eight participants, the site-disjoint residual differed by site (Kruskal–Wallis $H(16) = 50.7$, $p < 0.001$), but sites are nested in folds in this design and the residual is not norm-normalized, so this statistic confounds site with fold-model scale; among sites sharing a fold, a site effect was present in one of five folds ($p = 0.029$; others $p \geq 0.09$).

The pooled site-disjoint and stratified-split estimates were compared with a paired participant bootstrap (2000 draws) in which each draw resampled the same participants for both residuals: the difference was $-0.06$ (95% CI $-0.14$ to 0.02, $p = 0.12$). Each configuration was trained once with the same seeds, so this interval reflects participant sampling and not variability between retrains.

In the prospective and site-disjoint models reported earlier, early stopping and checkpoint selection used the morphometric loss on the held-out fold. Both models were retrained with an inner validation set of 15% of each fold's real training participants (stratified by diagnosis, seed 42 plus the fold index), excluded from gradient updates and used for early stopping and checkpoint selection; the held-out fold was scored only after training. Group separation was $H(2) = 16.1$ ($p < 0.001$, $\varepsilon^2 = 0.053$) in the stratified split and 14.2 ($p < 0.001$, $\varepsilon^2 = 0.047$) in the site-disjoint folds, compared with 19.1 and 19.0 with held-out-fold stopping. The partial association with 24-month MMSE change was $\rho = -0.15$ ($p = 0.009$) and $-0.18$ ($p = 0.001$), compared with $-0.14$ and $-0.20$. Within AD it was $\rho = -0.08$ ($p = 0.47$) and $-0.16$ ($p = 0.14$), compared with $-0.12$ and $-0.21$. Held-out-fold stopping therefore did not account for the group separation or the pooled association; group separation was smaller under inner-validation stopping, which also trains on 15% fewer real participants.

MMSE change from baseline to month 24 overlaps the month-12 imaging input of the prospective models. Change from month 12 to month 24, which follows all imaging inputs, was therefore analysed with month-12 MMSE taken from the ADNI MMSE table and matched by participant identifier ($n = 303$; mean change $-1.2$ points pooled and $-2.4$ points in AD). Controlling for atrophy and month-12 MMSE, neither prospective model was associated with this change (Supplementary Table S1).

**Table S1. Association with MMSE change from month 12 to month 24.** Partial Spearman correlation between the prospective residual and month-12-to-month-24 MMSE change, controlling for month-24 mean atrophy and month-12 MMSE (participant bootstrap 95% CI, 2000 draws). The baseline-to-month-24 estimates are shown for reference.

| Model | Outcome interval | Scope ($n$) | Partial $\rho$ (95% CI) | $p$ |
|---|---|---|---|---|
| Stratified split | baseline to month 24 | pooled (303) | −0.14 | 0.017 |
| Stratified split | month 12 to month 24 | pooled (303) | −0.05 (−0.16, 0.07) | 0.42 |
| Stratified split | month 12 to month 24 | AD (89) | −0.16 (−0.38, 0.07) | 0.13 |
| Site-disjoint | baseline to month 24 | pooled (303) | −0.20 | < 0.001 |
| Site-disjoint | month 12 to month 24 | pooled (303) | −0.04 (−0.15, 0.06) | 0.46 |
| Site-disjoint | month 12 to month 24 | AD (89) | −0.16 (−0.36, 0.06) | 0.14 |

#### Supplementary Note 5. Auxiliary heads and rate adjustment

A logistic regression on the two input features that define the tipping target (within-subject volume slope and month-12 volume) achieved per-region AUCs of 0.991 to 0.996 at month 12 without graph, attention, or temporal encoding (0.989 to 0.994 under five-fold cross-validation), and the trained head, evaluated in held-out participants of the three-visit fold models, achieved 0.998 to 1.000; the head therefore learns a near-deterministic rule, and its output is excluded. The no-tipping residual was strongly correlated with the reported residual (Pearson $r = 0.89$); its group contrasts were AD versus CN $p = 0.010$, AD versus MCI $p = 0.001$, and CN versus MCI $p = 0.89$, and its group separation was below the range of the other configurations ($H = 18.4$ to 21.8). In the no-tipping model, cross-validated ridge regressions predicting the month-12 rate and month-24 volume from the embedding gave $R^2 = 0.94$ and 0.83, respectively. With all five regional rates added to the covariates, the association was $\rho = -0.12$ ($p = 0.036$) in the stratified split, $\rho = -0.18$ ($p = 0.002$) in the site-disjoint model, and $\rho = -0.14$ ($p = 0.016$) in the no-tipping model; in the stratified atrophy-loss-only model it remained near zero ($\rho = -0.03$, $p = 0.66$).

#### Supplementary Note 6. Predictive uncertainty, gradient attribution, and representational similarity

For each held-out participant we ran 50 stochastic forward passes with dropout active [1] (the architecture uses layer normalization only, so no running statistics are affected) and took the variance of the atrophy-head prediction as predictive uncertainty. Uncertainty separated the diagnostic groups ($H = 9.87$, $p = 0.007$), correlated with the residual (Spearman $\rho = 0.50$, $p < 0.001$), and was associated with MMSE decline (partial $\rho = -0.15$, $p = 0.010$). Because both quantities reflect local instability of the same embedding, this agreement is supportive but not independent evidence. Integrated Gradients [2] was computed with 50 integration steps, the summed atrophy-head output as target, and each participant's diagnostic-group mean as the reference input. Mean absolute attribution was approximately uniform across regions (0.097 to 0.106) and greatest for month 24, followed by month 12 and baseline (0.146, 0.101, and 0.057). This analysis describes the sensitivity of the atrophy-head output, not changes in attention across visits. In a representational-similarity analysis of the held-out three-visit embeddings (cosine distance), pairwise embedding distance was unrelated to pairwise differences in month-24 atrophy when pairs were pooled across fold models (Mantel $\rho = 0.004$), because independently trained fold models do not share a coordinate frame. Within fold models, the association was strong ($\rho = 0.38$, within-fold label permutation $p < 0.001$), as expected for a model that receives month-24 morphometry, and weak for diagnosis ($\rho = 0.07$, $p < 0.001$) and *APOE* $\varepsilon 4$ carrier status ($\rho = 0.03$, $p = 0.006$). Downstream analyses therefore used residuals computed within fold rather than pooled embedding geometry.

#### Supplementary Note 7. Trial-enrichment analysis

AD participants were split at the median residual, and MMSE change was compared between strata with a Mann–Whitney test and Cohen's $d$. Treating stratum-specific means and standard deviations as trial parameters, we computed the per-arm sample size for a two-sided two-sample test ($\alpha = 0.05$, 80% power) and bootstrapped it by resampling AD participants (2000 draws). In the prospective model, mean MMSE change was $-5.2$ versus $-4.0$ points in the high- and low-residual strata, the 95% CI of the per-arm sample-size reduction spanned $-25\%$ to $+46\%$, and an upper-tertile split was also not significant ($d = -0.15$, $p = 0.46$). Analyses use the 89 AD participants with month-24 MMSE. For coarse enrichment after adjustment for atrophy, the residual performed comparably to the Mahalanobis and normative scores.

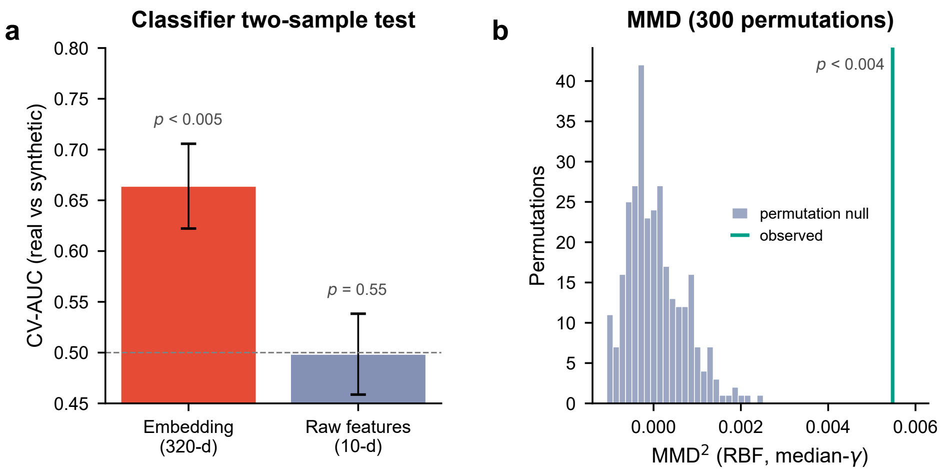


**Figure S1. Synthetic-data audit of the embedding.** Three-visit fold-4 model (lowest validation loss), applied to the 305 real and 1000 synthetic participants of the three-visit training set. **a**, Five-fold cross-validated AUC of a logistic classifier separating real from synthetic participants in the 320-dimensional embedding and in a ten-dimensional baseline volume-and-thickness feature set (error bars, standard deviation across folds; $p$ from 200 label permutations, reported at the permutation resolution). **b**, Squared maximum mean discrepancy (RBF kernel, median-heuristic bandwidth) between real and synthetic embeddings: permutation null (300 permutations) and observed value.

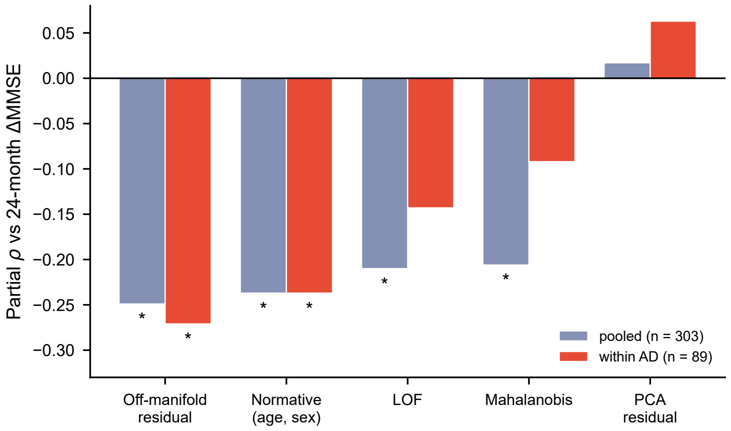


**Figure S2. Comparison with conventional anomaly scores.** Partial Spearman correlation between each score and concurrent 24-month MMSE change, controlling for atrophy and baseline MMSE, pooled and within AD. All scores use the same input features (five regions × seven features × three visits) and held-out participants as the three-visit model. Asterisks, nominal $p < 0.05$ for the individual association; they do not denote a significant difference between scores.

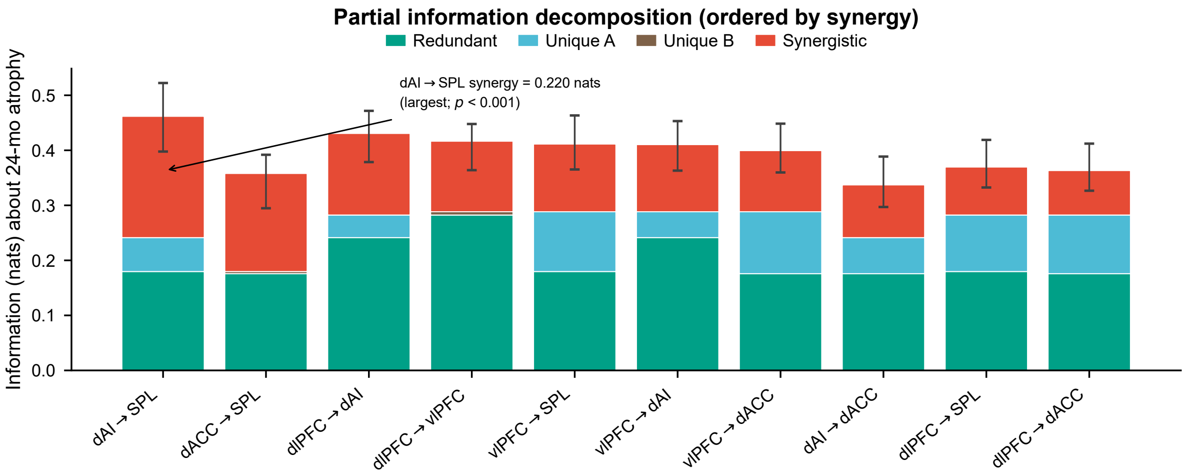


**Figure S3. Partial information decomposition over region pairs.** Gaussian-copula partial information decomposition of the information each region pair carries about month-24 atrophy. Sources are baseline regional volumes and the target is the mean month-24 volume ($n = 305$). The dAI–SPL pair carries the largest synergy (0.220 nats; one-sided permutation $p < 0.001$; percentile bootstrap 95% CI 0.156 to 0.281 nats); prefrontal executive-control pairs are dominated by redundancy.

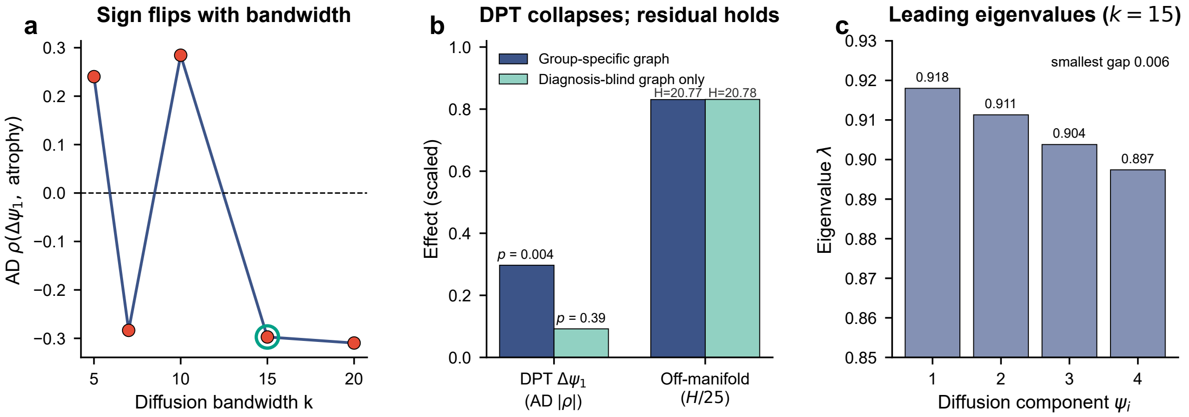


**Figure S4. Diffusion pseudo-time control.** **a**, Spearman correlation between $\Delta\psi_1$ and month-24 atrophy within AD as a function of diffusion bandwidth; significant values of both signs occur (red, $p < 0.05$), and the default setting ($k = 15$, green ring) is not representative. **b**, The same correlation in the reported model and in the graph-only diagnosis-blind retrain (pooled edges, diagnostic metadata retained), alongside the off-manifold residual group statistic ($H/25$; $H = 20.77$ and 20.78, two separately trained models whose values agree to one decimal place); this graph-only control is distinct from the fully diagnosis-blind retrain ($H = 21.5$). **c**, The four leading non-trivial diffusion eigenvalues at the default bandwidth ($k = 15$; smallest leading eigengap 0.006).

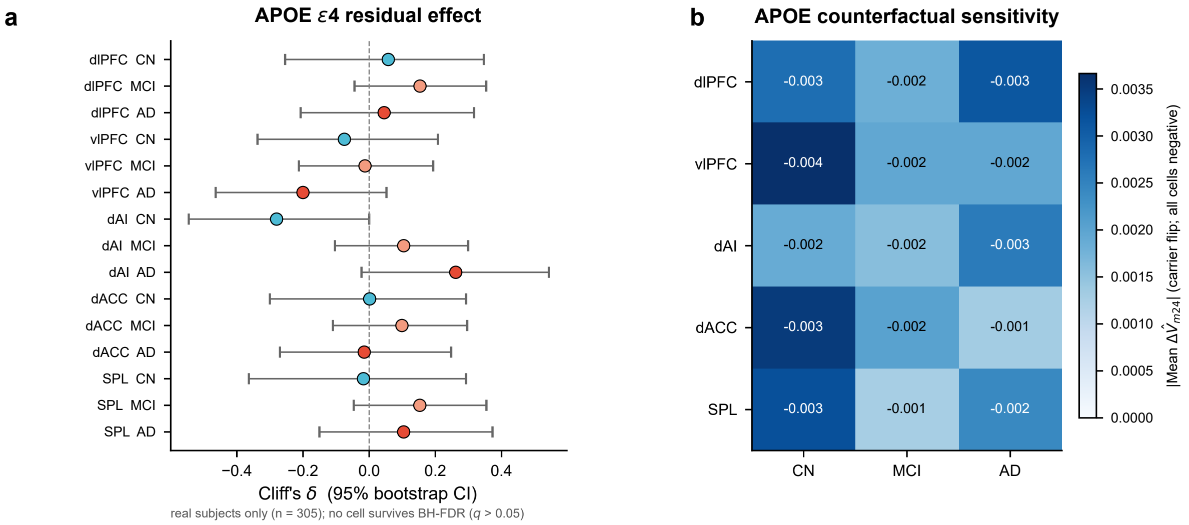


**Figure S5. *APOE* ε4 effects on prediction residuals.** Each participant was evaluated by the three-visit fold model that was not trained on them. **a**, Cliff's delta [3] of carrier-minus-non-carrier month-24 prediction residuals by region and diagnostic group, with percentile bootstrap 95% CIs [4]; no cell survives Benjamini–Hochberg correction over the 15 cells (smallest $q = 0.39$). **b**, Mean change in predicted month-24 volume, oriented towards carrier status, after setting each non-carrier's genotype inputs to $\varepsilon3/\varepsilon4$ and each carrier's to $\varepsilon3/\varepsilon3$ and re-running the model, by region and diagnostic group. All 15 cell means are negative; labels show signed means and colour shows their magnitude.

**Table S2. Cognitive endpoints.** Partial Spearman correlation between the off-manifold residual and 24-month change in each ADNI neuropsychological measure, pooled, controlling for atrophy and the measure's baseline score. $q$ is Benjamini–Hochberg-corrected across the six endpoints for each model; no endpoint hierarchy was pre-specified. Baseline logical memory is the screening-visit score. Scores recorded with an ADNI missing-value code at either visit were treated as missing, so $n$ differs by endpoint. Trail Making Test B is timed, so decline is a positive change; for the other measures decline is a negative change. Only MMSE change reached the corrected threshold, in the three-visit and site-disjoint models. Within AD, no endpoint reached it in any model (MMSE: three-visit $q = 0.067$, prospective $q = 0.69$, site-disjoint $q = 0.32$).

| Endpoint | $n$ | $\rho_{\text{partial}}$ | raw $p$ | $q$ |
|---|---|---|---|---|
| *Three-visit model* | | | | |
| MMSE (MMSCORE) | 303 | −0.249 | < 0.001 | < 0.001 |
| Boston Naming Test (BNTTOTAL) | 294 | −0.133 | 0.023 | 0.070 |
| Animal fluency (CATANIMSC) | 298 | −0.102 | 0.080 | 0.161 |
| RAVLT delayed recall (AVDELTOT) | 289 | −0.019 | 0.746 | 0.830 |
| Trail Making Test B (TRABSCOR) | 269 | 0.077 | 0.212 | 0.318 |
| Logical memory delayed (LDELTOTAL) | 294 | 0.013 | 0.830 | 0.830 |
| *Prospective model* | | | | |
| MMSE (MMSCORE) | 303 | −0.138 | 0.017 | 0.100 |
| Boston Naming Test (BNTTOTAL) | 294 | −0.088 | 0.135 | 0.258 |
| Animal fluency (CATANIMSC) | 298 | −0.080 | 0.172 | 0.258 |
| RAVLT delayed recall (AVDELTOT) | 289 | 0.013 | 0.830 | 0.830 |
| Trail Making Test B (TRABSCOR) | 269 | 0.045 | 0.461 | 0.554 |
| Logical memory delayed (LDELTOTAL) | 294 | 0.090 | 0.126 | 0.258 |
| *Site-disjoint model* | | | | |
| MMSE (MMSCORE) | 303 | −0.200 | < 0.001 | 0.003 |
| Boston Naming Test (BNTTOTAL) | 294 | −0.100 | 0.090 | 0.220 |
| Animal fluency (CATANIMSC) | 298 | −0.093 | 0.110 | 0.220 |
| RAVLT delayed recall (AVDELTOT) | 289 | −0.026 | 0.665 | 0.798 |
| Trail Making Test B (TRABSCOR) | 269 | 0.062 | 0.315 | 0.472 |
| Logical memory delayed (LDELTOTAL) | 294 | 0.000 | 0.999 | 0.999 |

**Table S3. Selected inferential results.** Significance is judged on the Benjamini–Hochberg $q$ where a correction is reported and otherwise on the raw $p$, at 0.05. MMSE and BNT associations are partial Spearman correlations adjusted for the covariates in Methods, with $q$ corrected across the six cognitive endpoints; enrichment $p$ values are from Mann–Whitney tests, with Cohen's $d$ as effect size. Dashes indicate values not shown. Site-disjoint, site-adjusted, rate-adjusted, and ablation estimates are shown in Fig. 4.

| Test | Model | Scope | Statistic | raw $p$ | $q_{\mathrm{BH}}$ | Significant |
|---|---|---|---|---|---|---|
| Group separation | three-visit | CN/MCI/AD | $H = 20.8$ | $< 0.001$ | — | yes |
| Residual–atrophy association | three-visit | pooled | $\rho = 0.12$ | 0.035 | 0.14 | no |
| Residual→MMSE | three-visit | pooled | $\rho = -0.25$ | $< 0.001$ | $< 0.001$ | yes |
| Residual→MMSE | three-visit | AD | $\rho = -0.27$ | 0.011 | 0.067 | no |
| Residual→BNT | three-visit | pooled | $\rho = -0.13$ | 0.023 | 0.070 | no |
| Group separation | prospective | CN/MCI/AD | $H = 19.1$ | $< 0.001$ | — | yes |
| Residual→MMSE | prospective | pooled | $\rho = -0.14$ | 0.017 | 0.100 | no |
| Residual→MMSE | prospective | AD | $\rho = -0.12$ | 0.270 | 0.690 | no |
| Residual→MMSE | site-disjoint | pooled | $\rho = -0.20$ | $< 0.001$ | 0.003 | yes |
| Residual→MMSE | site-disjoint | AD | $\rho = -0.21$ | 0.053 | 0.316 | no |
| Enrichment (median split) | three-visit | AD | $d = -0.41$ | 0.046 | — | yes |
| Enrichment (median split) | prospective | AD | $d = -0.21$ | 0.400 | — | no |
| Attention dAI→SPL | three-visit | pooled | $\rho = -0.19$ | — | 0.005 | yes |
| Pseudo-time (control) | diagnosis-blind | AD | $\rho = 0.09$ | 0.390 | — | no |

### Supplementary references

[1] Yarin Gal and Zoubin Ghahramani. Dropout as a Bayesian approximation: Representing model uncertainty in deep learning. In *International Conference on Machine Learning*, 2016.

[2] Mukund Sundararajan, Ankur Taly, and Qiqi Yan. Axiomatic attribution for deep networks. In *International Conference on Machine Learning*, 2017.

[3] Norman Cliff. Dominance statistics: Ordinal analyses to answer ordinal questions. *Psychological Bulletin*, 114:494–509, 1993.

[4] Bradley Efron. Better bootstrap confidence intervals. *Journal of the American Statistical Association*, 82:171–185, 1987.
