## Supplementary material for "A Candidate Trajectory-Atypicality Marker of Concurrent Cognitive Change in Alzheimer’s Disease from a Spatio-Temporal Graph Attention Network": TRIPOD+AI reporting checklist

Items follow the TRIPOD+AI statement (Collins et al., *BMJ* 2024;385:e078378); item wording is abbreviated. The study develops and internally validates a marker derived from a learned representation rather than a deployed clinical prediction model, so some items apply only in part. "Not reported" and "Not done" are stated where they apply. Locations refer to the main text unless marked SI (Supplementary Information).

| Item | Topic | Where reported / comment |
|---|---|---|
| 1 | Title: study type and target population | Title (candidate marker; Alzheimer's disease). |
| 2 | Abstract | Abstract. |
| 3a | Background: healthcare context and rationale | Introduction, paragraphs 1–4. |
| 3b | Target population and intended purpose | Introduction, paragraphs 1 and 4; Conclusion (candidate marker, not for clinical use). |
| 3c | Known health inequalities | Not reported. ADNI-1 is a volunteer research cohort; representativeness is not analysed. |
| 4 | Objectives (development, evaluation, or both) | Introduction, final paragraph (development with internal and site-disjoint validation). |
| 5a | Data sources | Methods, Participants (ADNI-1). |
| 5b | Dates of data collection | Not reported. |
| 6a | Setting and number of centres | Methods, Site-disjoint validation (47 acquisition sites). |
| 6b | Eligibility criteria | Methods, Participants. |
| 6c | Treatments received | Not applicable (observational cohort; treatments not analysed). |
| 7 | Data preparation and quality control | Methods, Feature extraction; Participants (quality-control exclusions). |
| 8a | Outcome definition and timing | Methods, Cognitive endpoints (24-month MMSE change, primary; five secondary endpoints). |
| 8b | Outcome assessment and blinding | Outcomes are ADNI clinical assessments made without knowledge of the marker; MMSE was not a training target (Discussion, paragraph 1). |
| 8c | Number of outcome assessors | Not applicable (standardized instrument scores from ADNI). |
| 9a | Predictors: definition and timing | Methods, Feature extraction; Off-manifold residual. |
| 9b | Predictor assessment and blinding | Automated image processing; Methods, Feature extraction. |
| 9c | Number of predictor assessors | Not applicable (automated). |
| 10 | Sample size | Methods, Participants: no formal calculation; all eligible participants analysed ($n = 305$). |
| 11 | Missing data | Methods, Participants (exclusion above 25% missing features); Cognitive endpoints (missing-value codes in secondary endpoints). |
| 12a | Use of data: partitioning | Methods, Model architecture and training (five-fold cross-validation); Held-out-fold protocol; Site-disjoint validation. |
| 12b | Handling of predictors | Methods, Feature extraction (features, z-scoring); Leakage controls (pooled standardization disclosed). |
| 12c | Model type, building, internal validation | Methods, Model architecture and training; Model variants and validation protocol; Table 2. |
| 12d | Heterogeneity across clusters | Methods, Site-disjoint validation (site omission, site indicators, site-clustered bootstrap). |
| 12e | Performance measures | Methods, Performance metrics; Statistical analysis. |
| 12f | Model updating / recalibration | Not done. |
| 12g | Evaluation procedures for model variants | Methods, Leakage controls; Auxiliary-head ablations; Comparison analyses and controls. |

| Item | Topic | Where reported / comment |
|---|---|---|
| 13 | Class imbalance | Folds stratified by diagnosis (Methods, Model architecture and training); no resampling. Synthetic augmentation: Methods, Synthetic augmentation and audit. |
| 14 | Fairness | Association of the marker with age, sex, and *APOE* $\varepsilon 4$ reported (Results, Fig. 2c); no subgroup performance analysis. Race, ethnicity, and education not analysed (Limitations). |
| 15 | Model output | Methods, Off-manifold residual (continuous score; no decision threshold proposed). |
| 16 | Differences between development and evaluation data | Same cohort; site-disjoint folds (Methods). No external evaluation (Limitations). |
| 17 | Ethical approval and consent | Ethics statement. |
| 18a | Funding | Funding. |
| 18b | Conflicts of interest | Competing interests. |
| 18c | Protocol | No study protocol was prepared (Methods, Cognitive endpoints: endpoint hierarchy is an a priori analytic decision). |
| 18d | Registration | Not registered (Methods, Cognitive endpoints). |
| 18e | Data sharing | Data and code availability (ADNI data-use agreement). |
| 18f | Code sharing | Data and code availability (on request). |
| 19 | Patient and public involvement | None. |
| 20a | Participant flow | Methods, Participants (criteria and final $n$); numbers excluded at each step not reported. |
| 20b | Participant characteristics | Table 1. |
| 20c | Comparison of development and evaluation data | Not applicable (cross-validation within one cohort). |
| 21 | Participants and outcome events in analyses | Table 1; Table 3; SI Table S1 (per-endpoint $n$). |
| 22 | Full model specification | Methods, Model architecture and training; Fig. 1; Software and reproducibility. Weights not public (Data and code availability). |
| 23a | Model performance with confidence intervals | Results; Table 3; Figs. 2–4. |
| 23b | Heterogeneity in performance | Results, site-disjoint validation; Fig. 4; SI Note 4. |
| 24 | Results of model updating | Not applicable. |
| 25 | Interpretation | Discussion, paragraphs 1–5. |
| 26 | Limitations | Discussion, limitations paragraph and falsifiability paragraph. |
| 27a | Handling of poor-quality or unavailable input | Not addressed; scans failing quality control were excluded (Methods, Participants). |
| 27b | User interaction and expertise required | Not applicable (no deployment proposed). |
| 27c | Next steps and future research | Discussion, final paragraph; Conclusion. |
